# *In silico* engineered multitarget-directed ligands for the polypharmaceutical treatment of PTEN loss of function endometrial adenocarcinoma

**DOI:** 10.64898/2026.08.28.747865

**Authors:** Ritchie Delara, Vaidehi Mujumdar, Qi Zhang, Hailey Dryden, Erin Crane, Jubilee Brown, Wendel Naumann, Allison Puechl, David Foureau, Wei Sha, Jessica LeGrand, Hsih Te Yang, Karl Dykema, Bharath Yada, Cody C. McHale, Krishnaiah Maddeboina, Dhananjaya Pal, Donald L. Durden

## Abstract

To combat refractory diseases, such as cancer, multitarget-directed ligands (MTDLs) have become an emerging area of research to exploit synthetic lethality (SL) relationships associated with drug resistance. Herein, we present the *in silico* design of MTDLs for the polypharmaceutical treatment of endometrial adenocarcinoma (EAC) and our discovery of a novel SL in EAC; PTEN loss of function (LOF) and the inhibition of CDK9. We used high-resolution x-ray crystallographic data to chemically engineer, LCI133, to inhibit CDK9, CDK4/6-and AURKA/B kinases. PTEN LOF in EAC results in augmented deregulated transcription and a massive increase in nascent RNA, a phenotype which encodes a high level of apoptotic sensitivity to LCI133 and CDK9 inhibitors. Treatment with LCI133 results in a rapid decline “**nose-dive**” in global nRNA, MYC nRNA levels and TS elongation (TE) in PTEN LOF EAC. PTEN LOF is *necessary and sufficient* to confer sensitivity of EAC cells to LCI133 and other CDK9 inhibitors.

**Statement of significance:** To exploit a novel pharmacologic SL, we *in silico* engineered the highly selective triple-action small molecule CDK9-CDK4/6-AURKA/B inhibitor, LCI133 which induces cancer cell death selectively in PTEN LOF EAC cells, an effect causally linked to PTEN loss of function, CDK9 inhibition and its associated deregulated/augmented TS elongation phenotype, transcriptional addiction (TA).

**Graphical abstract:** 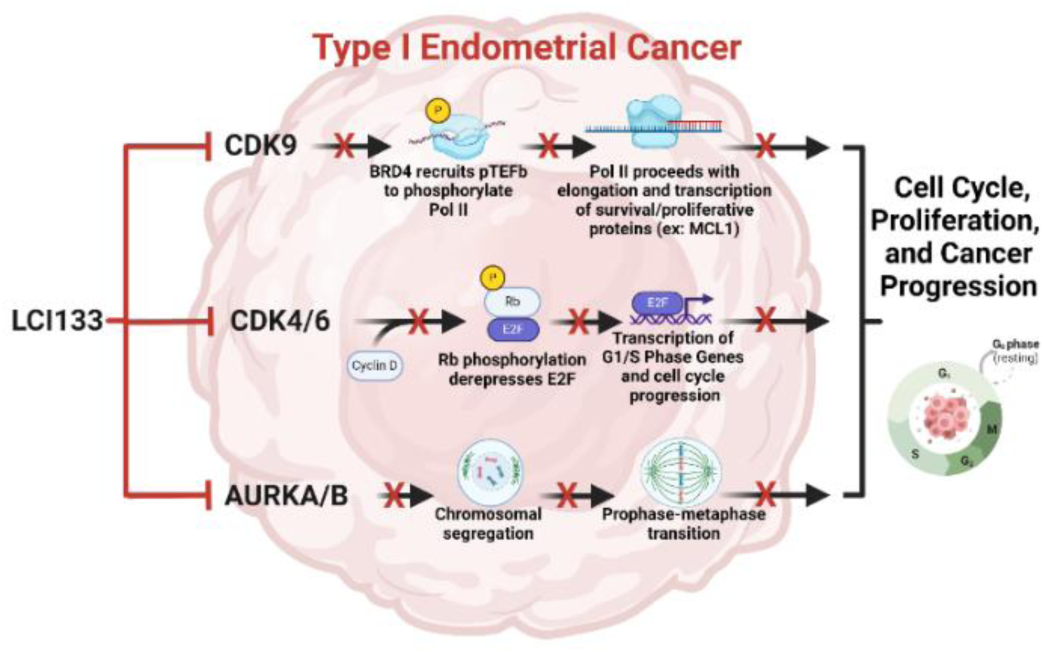

Schematic diagram shows the rationale for the intentional *in silico* design of the triple inhibitory chemotype, LCI133 to block three selective clinically relevant targets in PTEN mutated EAC to *in vivo* perturb the cell cycle, transcription (TS) and mitosis in same cancer cell at the same time (SL).

## Introduction

As treatment of cancer with a single targeted therapeutic agent, invariably results in tumor cell resistance and relapse of disease, we were interested in the intentional *in silico* design of one small molecule which binds selectively with nM potency to three different synergistic target molecules, CDK9, CDK4/6 and Aurora kinase A/B. The rationale for this polypharmaceutical approach comes from a number of reports that demonstrate: **1**) the difficulty in the treatment of cancer patients with 2 or 3 separate targeted therapeutic agents as result of toxicity and **2**) reports that treatment with dual or triple inhibitory chemotypes is safer and more efficacious compared to the high level of toxicity associated with treatment with 2 or 3 separate agents against the same targets with equal potency ^1–3^. The failure of single agent targeted therapy combined with these observations has stimulated considerable interest in the <u>intentional</u> (not by serendipity or high-through-put screens) *in silico* design of multitarget-directed ligands (MTDL) against specific targets for refractory cancers and to prevent the development of resistance ^4–7^.

Endometrial adenocarcinoma (EAC) is the most common type of uterine cancer and the most common gynecologic malignancy in the United States and is one of few cancers with increasing incidence and mortality ^8^. It is estimated that in 2024, nearly 68,000 women will be diagnosed with endometrial cancer (EC) and 13,000 will die of the disease in the United States. Few treatment options are available for advanced disease where the 5-year relative survival rate is 18%. The recent integrated genomic characterization of endometrial cancers by The Cancer Genome Atlas (TCGA) research network, created four prognostic subgroups: **1**) POLE ultramutated (good prognosis), **2**) high microsatellite instability (MSI-H)/defective mismatch repair (dMMR), **3**) TP53 wild type copy number low/non-specific molecular profile (NSMP) and **4**) TP53 mutated copy number high (poor prognosis) ^9^. While the TCGA was instrumental in the genomic and molecular characterization of EAC and the identification of potential actionable mutations, the application of targeted therapeutics remains limited ^10^.

The phosphatase and tensin homolog (PTEN) is a dual specificity phosphatase, tumor suppressor gene that is mutated in 85% of EAC cases. Within TCGA EAC prognostic subgroups, PTEN alterations occur at 87.5% (POLE), 75% (MSI-H/dMMR), 60.4% (CNL), and 5% (CNH). PTEN mutations occur at high frequency in 3 of the 4 TCGA classification groups and occur in endometrioid low-grade and high-grade high-risk EC within the POLE, MSI and CNL groups of EC which have 75% long term DFS ^9^. It is commonly accepted that PTEN mutations occur early in EAC oncogenesis and co-occur with other alterations to the phosphatidylinositol-3-OH kinase PI3K/protein kinase B (AKT)/mammalian target of rapamycin (mTOR) pathways ^10^. The majority of PTEN alterations in EAC result in loss of function (LOF) point mutations or deletions, driving the buildup of PIP3 at the plasma membrane and activating PI3K/AKT/mTOR pathway-driven cell growth, proliferation, and survival ^11^. While PTEN is best known for its cytoplasmic role as a negative regulator of PI3K/AKT pathway signaling, nuclear PTEN plays a role in homologous recombination-mediated repair of double-strand DNA breaks, cell proliferation, chromatin condensation and the regulation of transcription (TS) ^12,13^. Despite the high frequency of PTEN loss of function and its canonical role in transcription, cell proliferation and survival in all subgroups of EAC, PTEN genomic status currently plays no role in disease risk stratification or therapy for this disease.

Cyclin dependent kinases (CDKs) are a family of 21 kinases involved in cell cycle progression (CDK4-6/14-18) and transcriptional regulation (CDK7, 8, 9-13/19-20) ^14^. CDK4/6 inhibitors (CDK4/6i), which block G1 to S cell cycle transition, have been FDA-approved for the treatment of hormone receptor-positive HER2-negative metastatic breast cancer since 2012. Additionally, the G2 to M mitotic phase of the cell cycle ending in cytokinesis is regulated by transcription under the control of CDK1/cyclin B1 and Aurora kinase A/B ^15^. Aurora kinase A and B are 71% homologous members of the mitotic kinome and are critical for orchestrating the initiation and overall process of mitosis. AURK A is involved in early stages of G2-M transition whereas AURK B functions later at metaphase to organize the spindle apparatus, microtubule alignment of chromosomes and cytokinesis. AURK A/B are overexpressed in a number of malignancies and are strongly associated with aneuploidy in most cancers including EAC where AURK B correlates with increases in Ki67 staining and predicts a poor prognosis^16^.

Finally, CDK9, the catalytic subunit of the positive transcriptional elongation factor b (P-TEFb) complex is responsible for RNA polymerase II (Pol II) pause release (PR) and Pol II-mediated productive elongation and termination of transcription (TS). P-TEFb/CDK9 has been described as a complex protein interaction network that is critical for the modulation of pro-proliferative, survival signals and coordinates the activity of many oncoproteins involved in the pathogenicity or maintenance of malignant phenotypes ^17^. CDK9 has recently been proposed as a biomarker and target for the treatment of EAC but little clinical progress has been made. In summary, the genomic landscape of endometrial cancer suggests it could be particularly susceptible to CDK9 or pan-CDK inhibition and/or inhibitors of transcription (TS)^14^. Finally, an analysis of a genome wide vulnerability CRISPR screen (DepMap) reveals CDK9 and CDK4 as target genes in EAC.

In our study, we chemically engineered three chemotypes, LCI132, LCI133 and LCI136 (**Figure 1**)^7^. These benzopyranone-based positional isomers underwent x-ray crystal structure-based carbon to nitrogen chemical modifications in order to modify the binding specificity and affinity toward each target protein e.g. PI3K, BRD4 or CDK4/6/9/Aurora kinase. LCI133 in cell free IC_50_ assays simultaneously inhibits CDK9 (4 nM), CDK4/6 (4.7/19.2 nM), AURK A/B (2.8/10 nM), is highly selective, displays favorable drug-like qualities, and is potent in *in vitro*, *in vivo* and in primary human tumor derived models of *PTEN*-mutant EAC. Furthermore, we discovered a role for PTEN LOF in the modulation of apoptotic sensitivity to LCI133 and CDK9i, a potential mechanistic biomarker which predicts an augmented and highly deregulated transcriptional elongation rate (TS addiction, TA) as a cause for EAC sensitivity to LCI133 and other CDK9 inhibitors (CDK9i).

**Figure 1.**
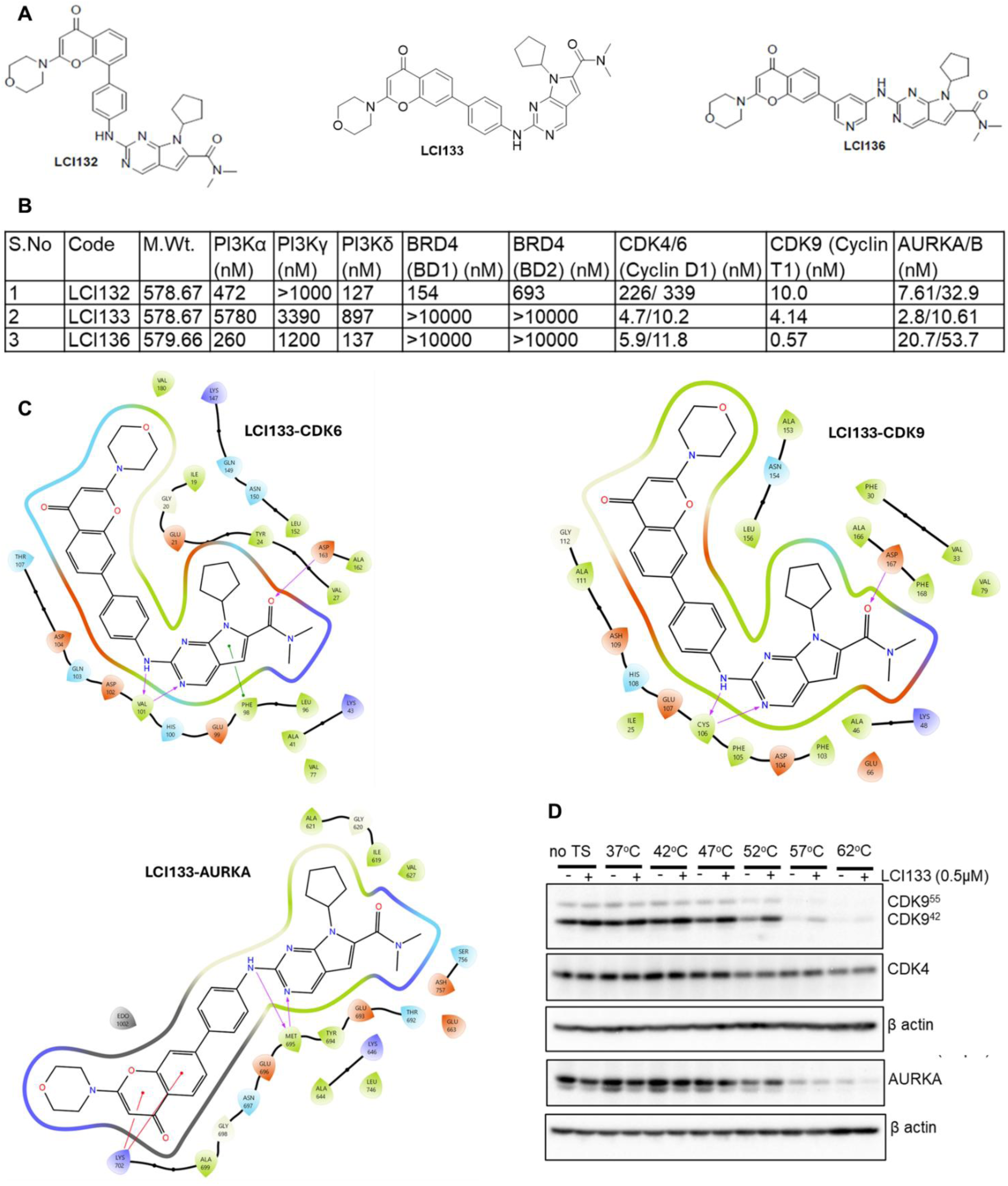
Molecular modeling results for LCI133 and potency results for LCI132, LCI133 and LCI136. **A,** Chemical structures of LCI132, LCI133, and LCI136. **B**, IC_50_ values (nM) of LCI132, LCI133 and LCI136 in cell free assays (**Fig S1** for raw datasets, IC50 curves). **C**, 2D representations of LCI133-CDK6, LCI133-CDK9, and LCI133-AURKA bound complexes, respectively. Pink color arrow indicates hydrogen bond, and green solid line represents predicted pi-pi stacking, and red solid line indicates pi-cation interaction. These computational data represent the fundamental and <u>only</u> method (***in silico*** design) and modeling used to design small molecules, to <u>selectively</u> bind to all targets at nM potency. **D**, Cellular thermal shift assay performed on RL95-2 cells treated with DMSO or LCI133 at 500 nM concentration and exposed to increasing temperatures for 20 minute time points followed by cell lysis, lysates were subjected to Western blot analysis to quantitate the relative stability of each target protein in the presence or absence of the small molecule ligand, LCI133 e.g. CDK9, CDK4, Aurora A kinase versus actin as control. Result were confirmed for K_d_ determinations via Nano-Bret analysis for each chemotype-target molecule interaction.

## Results

### *In silico* chemical engineering of LCI132, LCI133, LCI136 and cell free potencies

We used x-ray crystal structure datasets and Schrodinger software to engineer 3 chemotypes which bind to CDK4/6/9 and Aurora kinase plus or minus PI3K and/or BRD4^7^. The *in silico* design, synthesis and characterization of LCI132, LCI133, LCI136 and other chemotypes are described in detail elsewhere^7^. Cell free assays of potency against PI3K, CDK4/6, CDK9, AURK A/B and a high level of kinome selectivity for these targets was reported ^7^. The chemical structures for LCI132, LCI133 and LCI136 are shown in **Figure 1A**. In cell-free IC_50_ assays, LCI132 blocks PI3K-p110α, PI3Kδ, BRD4, CDK4/6, CDK9 and AURK A/B. LCI133 inhibits CDK4/6 at 4.7/10.2 nM, CDK9 at 4.14 nM and Aurora A/B kinases at 2.8/10.6 nM potency (**Figure 1B; raw data, S1**). Whereas LCI136 inhibits PI3Kα, PI3Kδ, CDK4/6, CDK9 and Aurora kinase A/B at nM potency as designed (**Figure 1, B**). We reported LCI133 is highly selective for these targets and when profiled against 468 kinases it inhibits 2.5 percent of the kinome and does not inhibit CDK1, CDK2, CDK3, CDK5, CDK7, CDK8 or CDK12 as determined in cell free assays screened at 1 uM concentration or in cell-free IC_50_ titrations ^7^. We determined the cell-free IC_50_ curves for LCI133 against CDK1/cyclin A, 6,960 nM; CDK2/cyclin E, 483 nM; CDK7/cyclin H, >10,000 nM and CDK12/cyclin K, 1,560 nM as reported (CDK11/cyclin L not available) (10)^7^. AZD4573 inhibits CDK1, CDK9 and not CDK7, 4, 6, or 2 ^18^. These results are confirmed using computational modeling of LCI133 in the active site of each of these target kinases. LCI132, LCI133 and 136 are benzopyranone-based positional isomers. LCI132, LCI133 and LCI136 underwent carbon to nitrogen chemical modification and represent our ongoing efforts to modify each compound based on crystal structure datasets to convert each triple inhibitor to a dual or single inhibitor as we have published^5^. Using a cellular thermal shift assay (CETSA) for CDK9, CDK4 and AURKA, a well-established method for evaluating drug-target binding that determines drug binding to its cognate targets in a cellular setting, we demonstrate the association between LCI133 and its intended target molecules. A series of CETSAs at various temperatures with DMSO or LCI133-treated endometrial cancer cell line, RL95-2, demonstrated that 52°C is a temperature at which CDK9 and AURKA protein stability increased and to a lesser extent in CDK4, providing further evidence that LCI133 binds to these three target proteins inside the EAC cell (**Figure 1D**). CETSA results were confirmed using Nano-Bret analysis of K_m_ for binding to each target molecule.

### Cytotoxicity of LCI132, 133, 136, AZD4573, BKM120 and Ribociclib in EAC cells

The IC_50_ cytotoxicity profile for seven different EAC cell lines was determined. Cell lines were stratified by PTEN mutational status including AN3 CA, RL95-2, Ishikawa, HCI-EC-23, HEC-1-A, HEC-1-B, and KLE. Cells were treated with increasing concentrations of LCI132, LCI133, LCI136 or single agent reference compound inhibitors of PI3K (BKM120), CDK4/6 (Ribociclib), CDK9 (AZD4573)(**Figure 2 B-I**) or BRD4 (PLX51107)(**Fig 2I**) for 48 hours (**Figure 2B-H**). Figure S2A summarizes the mutational status of PTEN and other PI3K/AKT pathway genes sourced from cell bank and Dependency Map (DepMap) Portal data (Broad Institute, Cambridge, MA), and Figure 2A shows the PTEN expression levels and p-AKT activation status by Western blot analysis in all EAC cell lines under study. As expected, the results confirmed the expression of PTEN in HEC-1-A, HEC-1-B, and KLE. PTEN was undetectable in AN3-CA, RL95-2 and Ishikawa cell lines. The dose-response curves in **Figure 2B-H** generated from Cell-Titer Glo cell viability assays show that while all three chemotypes displayed potency, LCI133 generated the most potent cytotoxic response with sub-500 nM IC_50_ of 249 nM in AN3 CA, 312 nM in RL95-2, 900 nM in Ishikawa and 23 nM in HCI-EC-23 EAC cells. Interestingly, the relative potency paralleled the potency of a reference compound CDK9i, AZD4573 against AN3 CA (.0024 uM), RL95-2 (.0027 uM), Ishikawa (.0019 uM) and HCI-EC-23 (.0236 uM). AN3 CA, RL95-2, and Ishikawa cell lines all possess *PTEN* frameshift mutations resulting in loss of detectable PTEN protein and HCI-EC-23 has a point mutation in the P-loop of PTEN (R130G) which neutralizes all PTEN phosphatase activity^19^. This was confirmed by Western blot analysis, PTEN loss of function (**Figure 2A**) cells displayed an increase in pAKT. These data suggested an association between PTEN loss of function and sensitivity to CDK9 inhibitory chemotypes. The pan-PI3K inhibitor BKM120 showed sub uM IC_50_ activity across PTEN-deleted and PTEN-wild type cell lines. While AZD4573 demonstrated higher potency (IC_50_ = 2 nM) in both AN3 CA and RL95-2, it also exhibited marked cytotoxicity in non-transformed HS5 human bone marrow stromal cells (**Figure S2C**), LCI133 demonstrated no detectable cytotoxicity in HS5 bone marrow stromal cells. To further validate these results, we performed colony formation assays by one-time treatment of AN3 CA or HEC-1-A cells with BKM120, 0.25 uM, PLX51107, 5.0 uM, AZD4573, 0.0005 uM, Ribociclib, 5.0 uM, LCI132, 5.0 uM, LCI133, 0.25 uM, and LCI136, 1 uM. Colony formation was assessed after a three-week incubation period. PLX51107, AZD4573, LCI133, and LCI136 decreased capacity of AN3 CA cells to form colonies (**Figure 2I**). HEC-1-A cells were most sensitive to PLX51107, Ribociclib, AZD4573, and LCI132, and moderately sensitive to LCI133 and LCI136. The augmented sensitivity to LCI133 in PTEN loss of function EAC cell lines was consistent with the results obtained from the quantitation of apoptosis (**Figure 3A-D**). LCI133 was nM potent against AN3 CA, RL95-2, Ishikawa and HCI-EC-23 but not HEC-1-A, HEC-1-B, and KLE **(Figure 2A-H)**. The direct correlation of PTEN LOF and IC_50_ potency of LCI133 and AZD4573 against EAC cell lines, suggested that PTEN mutational status may somehow determine sensitivity to LCI133 and/or CDK9 inhibitors (**Figure 2A-H**).

**Figure 2.**
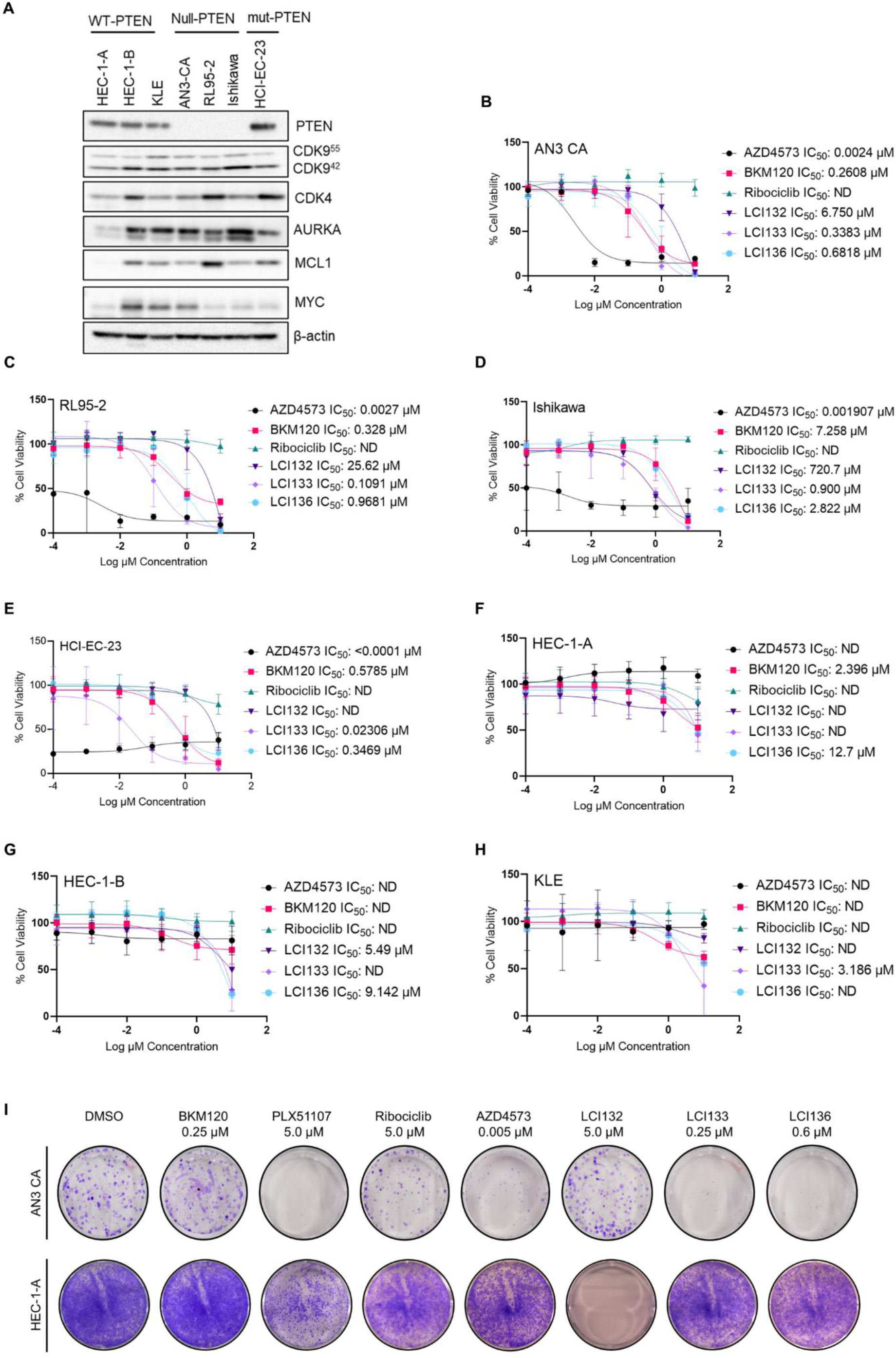
The novel BP scaffold multitarget inhibitor, LCI133 is differentially cytotoxic to PTEN mutant (Mut) versus PTEN wildtype (WT) EAC cells. **A**, Representative Western blot of EAC cell lines showing biochemical status of PTEN, LCI133 targets (CDK4, AURKA, CDK9), and pharmacodynamic markers of CDK9 inhibition (MCL-1, MYC) with β actin as loading control. **B-H**, PTEN-null AN3 CA (**B**) RL95-2 (**C**) Ishikawa (**D**) and HCI-EC-23 (**E**) and PTEN wild type HEC-1-A (**F**) HEC-1-B (**G**) and KLE **(H**) cell lines treated with increasing concentrations of the indicated inhibitors and evaluated 48-hours later for cell viability by CellTiter-Glo assay. Data represent the mean of triplicate experiments ± SD. (**I**) Representative Colony formation assay of AN3-CA (*top*) or HEC-1-A cells (*bottom*) treated with the indicated inhibitors and evaluated 21-days later by Crystal violet stain. Table of IC_50_ quantitation for 2B-H in **Fig. S2**.

**Figure 3.**
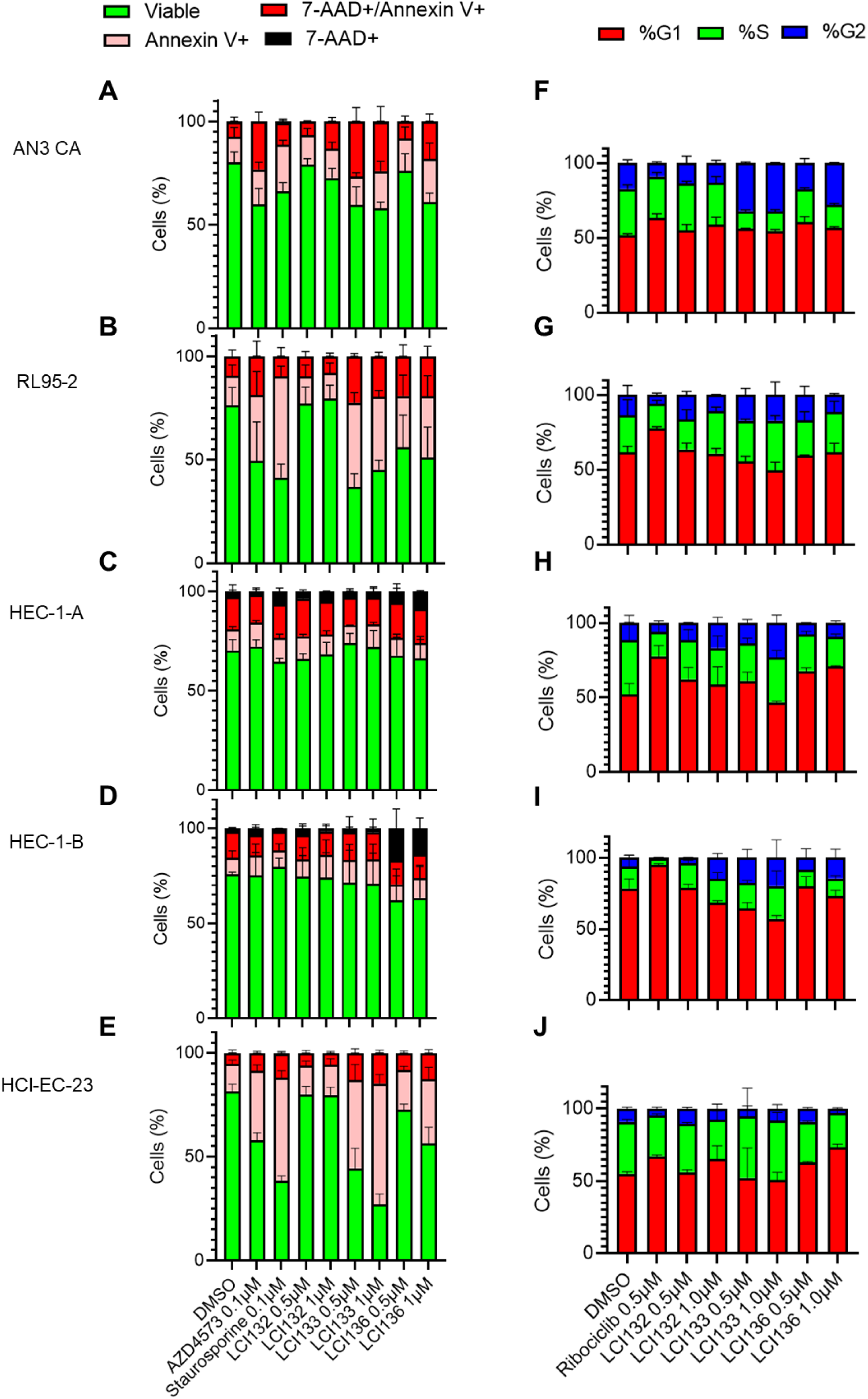
Differential apoptotic and cytostatic response of PTEN WT vs PTEN Mut EAC cell lines to single agent inhibitors and LCI132, LCI133, and LCI136. **A-E,** Flow cytometry analysis of apoptosis induced in AN3 CA (**A**) RL95-2 (**B**) HEC-1-A (**C**) HEC-1-B (**D**) and HCI-EC-23 (**E**) EAC cells treated with DMSO (-) control, staurosporine (+) control, or inhibitor at the indicated concentrations and assessed after 24-hours by evaluating the percentage of FITC-Annexin V/7-AAD stained cells.**F-J,** Flow cytometry analysis of cell cycle arrest induced in AN3 CA (**F**), RL95-2 (**G**) HEC-1-A (**H**) HEC-1-B (**I**), and HCI-EC-23 (**J**) EAC cells treated with DMSO (-) control, ribociclib (+) control, or inhibitors at the indicated concentrations and assessed after 24-hours by quantifying MFI distribution of DNA-binding dye. Data represent the mean of triplicate experiments ±SD. Quantitation of representative flow cytometry FACS analysis shown in **Fig. S3-4**. WB for PARP cleavage as an additional indicator of apoptosis shown in **Fig. S5.**

### LCI133 preferentially induces dose-dependent apoptosis in PTEN-mutated or loss of function EAC cells

Given the cytotoxicity results shown in Figure 2, we evaluated the effect of each chemotype on apoptosis. AN3 CA, RL95-2, HCI-EC-23, HEC-1A, HEC-1B cells were treated for 24 hours with LCI132, LCI133, LCI136 or single agent reference compounds, AZD4573 or staurosporine, at the indicated concentrations and cells were collected for assessment of apoptosis by flow cytometry and Western blot analysis (**Figure 3A-E and S3/S5**). The results demonstrate, while LCI132 (0.5 and 1.0 uM) had no significant effect on apoptosis in AN3 CA, HCI-EC-23, HEC-1A or HEC1B as measured by Annexin V/7-AAD staining, LCI133 (0.5 µM and 1.0 µM) and LCI136 (1.0 µM) induced dose-dependent marked increases in apoptosis in all PTEN mutated EAC cell lines but not in PTEN wild type EAC cells compared to DMSO at 24 hours (pFDR < 0.05) We generated confirmatory evidence in that 24-hour treatment with LCI133 (0.5 µM and 1.0 µM) or LCI136 (1.0 µM) resulted in a marked increase in the cleavage of poly (ADP-ribose) polymerase (PARP) to cPARP in all four PTEN mutant cell lines but not in PTEN wild type EAC cells compared to DMSO controls (pFDR < 0.05) (**Figure S5, A-E**). These data reveal increased apoptotic sensitivity to LCI133 and AZD4573 in PTEN mutated EAC cell lines compared to PTEN wild type EACs, (**Fig 3, A-E; S3, A-E, S5, A-E**). In contrast, we observed augmented cPARP in PTEN wild type HEC-1-A EAC cells treated with a BET inhibitor, PLX51107 compared to PTEN mutant EAC (**Figure S5, A-E**). Moreover, the marked increase in apoptosis as measured by cPARP Western blot was observed in response to LCI133 and AZD4573 in the PTEN mutated EAC cell line, HCI-EC-23 which suggests the loss of PTEN phosphatase activity is necessary and sufficient to directly drive apoptotic sensitivity to AZD4573 or LCI133 treatment (**Figure S5, C**).

### LCI133 induces dose-dependent G2 cell cycle arrest in all EAC cell lines

CDK4/6 inhibitors such as Palbociclib and Ribociclib induce G1 arrest by selectively binding to the ATP-binding pockets of CDK4-cyclin D3 and CDK6-cyclin D1 complexes with varying affinity, thus preventing the phosphorylation and inactivation of the retinoblastoma (Rb) protein and G1 to S phase transition. AURKA (e.g., Alisertib) and AURK B (e.g., Barasertib) inhibitors competitively bind to the ATP-binding pockets of their targets, delaying mitotic entry and progression from G2 to M phase. Interestingly, LCI133 and AZD4573 did not induce a G1 arrest as compared to positive control Ribociclib (**Figure 3, F-J, S4, A-D**). In contrast, LCI133 and LCI136 at 0.5 µM and 1.0 µM concentrations were found to significantly induce G2 arrest in all EAC cell lines compared to DMSO controls at 24 hours, with pFDR = 0.001 and 0.002, respectively (**Figure 3, F-J, S4, A-D**). Importantly, these data support the AURA/B inhibitory properties as LCI133 induced a similar level of G2 arrest in PTEN wild type and PTEN mutant EACs suggesting this is not the dominant mechanism for sensitivity to these chemotypes in PTEN loss of function EAC. AURK inhibition was confirmed by the examination effects of LCI133 on the phosphorylation status of histone (H3, S10), a known AURK substrate ^20^ in multiple EAC cell lines. Supporting this finding, while single agent AURKi, Alisertib similarly induced G2 cell cycle arrest *in vitro,* it was not cytotoxic to EAC cells (IC_50_: AN3 CA, 0.288 uM, RL95-2, ND or HEC-1-B, ND) where ND refers to a lack of sensitivity to this agent alone (data not shown).

### Effects if LCI133 and AZD4573 on canonical kinase and TS targets in PTEN mutant vs wild type EAC cell lines and on TS elongation and nRNA levels in PTEN mutant EAC cells

We next sought to compare the effects of LCI133 and AZD4573 on relevant kinase signaling pathways and on MYC/MCL-1 TS. To assess the effect of Ribociclib, AZD4573 or LCI133 on signaling pathways controlled by CDK4/6 and CDK9, we evaluated the effects on known canonical phosphorylation of specific protein substrates by CDK4/6, (pRb S780/811), and CDK9, (pRpb1 S2) (pPol II) (**Figure 4, A-D**). We determined the effects of LCI133 or AZD4573 on MYC and MCL1 protein and mRNA levels, both recognized canonical targets of CDK9 and AURK inhibition, respectively by Western blot and qRT PCR in EAC cell lines (**Figure 4, E-H**) ^21,22^. A 2 hour treatment of all EAC cell lines with LCI133 or AZD4573 inhibited the phosphorylation of the CDK9 target protein, Rpb1/Pol II (S2) and markedly decreased MCL-1 protein and mRNA levels in all EAC cell lines (**Figure 4, A-H**). LCI133 at 1.0 µM and AZD4573 at 100 nM significantly decreases MYC and MCL-1 gene expression after 3-hour and 6-hour treatments compared to DMSO (pFDR < 0.05) in AN3 CA cells (**Figure 4, E-H, S6B**). An increase in MYC gene and protein expression in AN3 CA was seen after 12 and 24 hours of treatment with LCI133 at 1.0 µM but not with AZD4573 at 0.1 uM (**Figure S6, C-E**). LCI133 and AZD4573 significantly decreases MCL1 gene expression in AN3 CA cells after 6-hour, 12-hour, and 24-hour treatments compared to DMSO (pFDR < 0.05) (**Figure S6, E**). This phenomenon has been reported by Lu et al. where they describe compensatory induction of MYC expression by sustained CDK9 inhibition via a BRD4-dependent mechanism ^23^. Others have reported that PTEN controls global gene transcription by the regulation of Pol II promoter-proximal pause release and may impact pause duration and release on MYC promoter transcriptional start sites (TSS)(+100 bp), elongation rates on gene bodies in a global manner ^13,24^. We examined the effects of LCI133 or the single agent CDK9i, AZD4573 treatment on MYC binding to its upstream proximal promoter and sites distal to the TSS (−350 bp) or downstream (+350-11,882) bp promoter gene body and termination zone using Pol II/pPol II CHiP qPCR methods in the RPL-95 PTEN mutant EAC cell line (**Figure 4,I**). The data demonstrate both LCI133 and AZD4573 block transcription of MYC and MCL1 mRNA in all EAC cell lines. Similarly, AZD4573 and LCI133 block Pol II occupancy of MCL1(not shown) and MYC promoters on the TSS (+100 bp), upstream and downstream gene body promoter regions (**Figure 4, I**). Our results show that AN3 CA cells display significant levels of nascent RNA (nRNA) by confocal microscopy of EU labeled nRNA (**Figure 4, J**). Treatment of these cells with AZD4573 or LCI133 led to a marked reduction in total nRNA levels in these cells (**Figure 4, J**) and the suppression of MYC and MCL nRNA bound to the MYC and MCL1 promoter regions (**Figure 4, K**).

**Figure 4.**
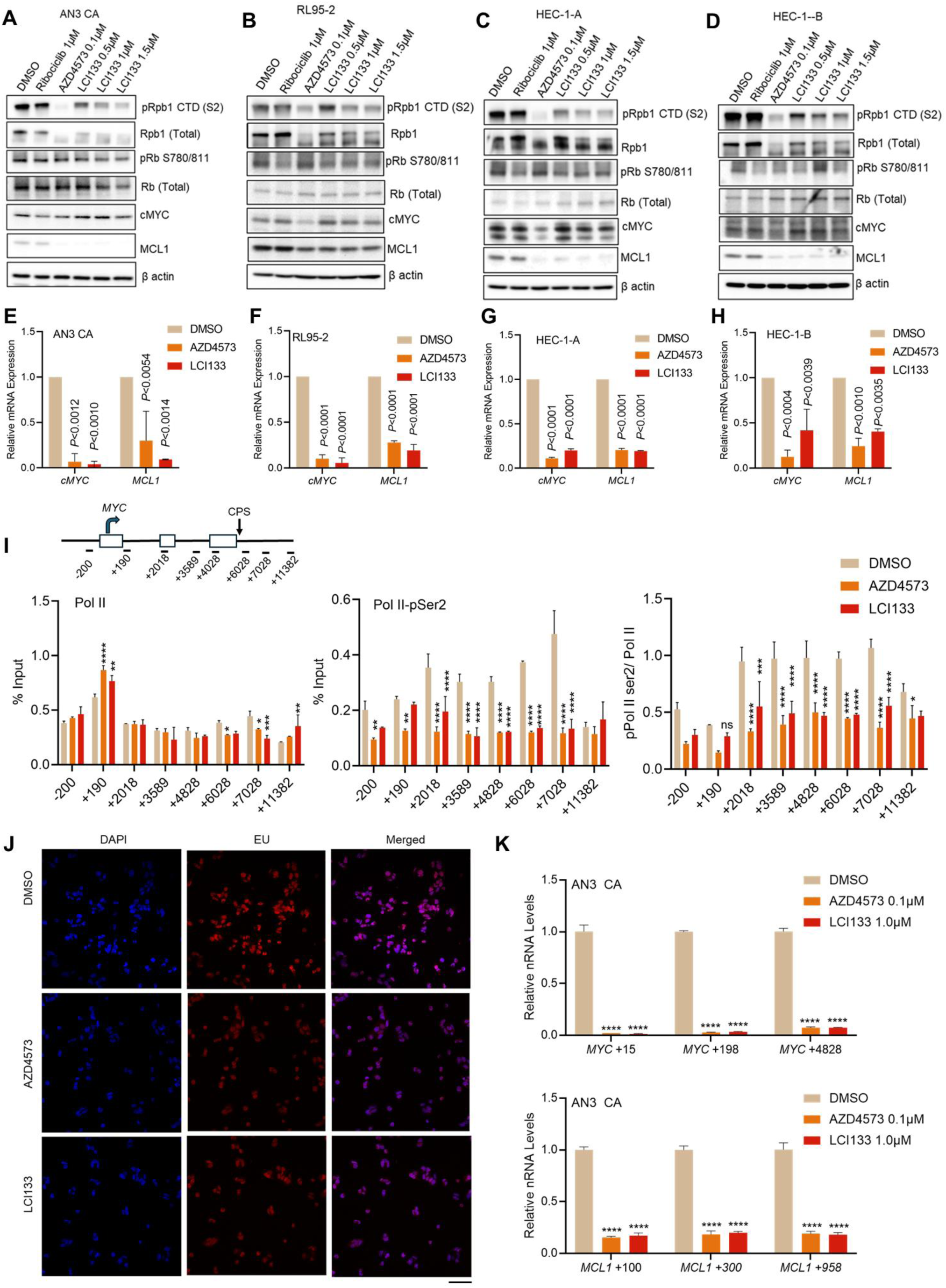
Effects of ribociclib, AZD4573, LCI133 on canonical biochemical target inhibition and AZD/ LCI133 on Pol II/pPol II occupancy of the MYC proximal and distal promoter regions total, MCL1/MYC specific a total nascent RNA (nRNA) levels in EAC cells. **A-D**, Representative immunoblots showing the effect of LCI133 and single agent positive control inhibitors AZD4573 (CDK9) ribociclib (CDK4/6) on RNA polymerase II (Rpb1), phosphorylated RNAP II at serine 2 in the C-terminal domain (p-Pol II CTD Ser 2), pRb at S780/T 811, total Rb, cMYC, and MCL1 in AN3 CA (**A**) RL95-2 (**B**) HEC-1-A (**C**) and HEC-1-B (**D**) EAC cell lines after 6-hour treatment. β actin was used as loading control. (**E-H**) Representative bar graphs of qRT-PCR data for expression of CDK9 target genes *MYC* and *MCL1* after 2-hour treatment with LCI133 or AZD4573 in AN3 CA (**E**) RL95-2 (**F**) HEC-1-A (**G**) and HEC-1-B (**H**) EAC cell lines. **I,** AN3 CA EAC cells were treated DMSO control, AZD4573 (0.1 μM), or LCI133 (1.0 μM) for 2-hours followed by Pol II ChIP qPCR analysis of *MYC* at eight different *MYC* promoter regions under study including TSS, gene body, and termination zone as represented in the schematic diagram. Data represent the mean ± S.E.M. biological duplicates. The numbers below the graph shows the position of primers from the TSS (bp). Data were analyzed using a two-way ANOVA. P value denotes a significant difference between treatment and vehicle, as determined using multiple comparisons test (as shown) or for Fig. 4I: ****p < .0001; ***p<0.001; **p < .01; and *p< 0.05. **J**, Representative confocal microscopy images showing effect of AZD4573 (0.1 μM) and LCI133 (1.0 μM) on the abundance of EU-labeled nascent RNA in AN3 CA cells compared to DMSO control. Scale bar: 10 µm. **K**, AN3 CA cells were treated with DMSO control, AZD4573 (0.1 μM), or LCI133 (1.0 μM) for 2-hours followed by EU-labelling and qRT-PCR assessment of *MYC* (upper) and *MCL1* (lower) nascent RNA. The y axis shows relative changes in nascent RNA levels normalized to total 7SK RNA between treated and untreated samples. Data are represented as mean ± S.E.M. from triplicates. ****p< 0.0001.

### PTEN is *necessary and sufficient* to regulate TS elongation, Pol II occupancy on the MYC promoter and global nRNA levels in EAC cells

In order to directly establish a causal role for PTEN in LCI133 and CDK9i EAC sensitivity and implicate PTEN in the regulation of EAC transcription, we assessed the potential protein-protein interaction between PTEN, Pol II and the TS apparatus. We demonstrated the reciprocal binding of PTEN to Pol II as well as cyclinT1, ELL2 and SPT5 in PTEN positive EAC cells (**Figure 5, A-D**). Moreover, PTEN shRNA KD in HEC-1-B and KLE cells was confirmed to induce a dose dependent suppression of PTEN, a corresponding increase in pAKT levels (**Figure S7,C, S8, A/B**) and a specific ablation of the co-immunoprecipitation of PTEN and components of the transcriptional mediator complex including components of the super elongation complex (SEC) including SPT5 (**Figure S6D**). Moreover, we demonstrate the KD of PTEN in HEC-1-B cells results in an increase in the ratio of chromatin associated pPol II/Pol II and pCTR1/CTR1 on the MYC promoter in EAC cells (**Fig. S8, C/D**). These data provide evidence that PTEN LOF drives an increase in pPol II and pCTR1 occupancy on chromatin i.e. the MYC promoter in EAC cells. We further demonstrated that PTEN shRNA in HEC-1-B and KLE cell lines resulted in increase of Pol II/pPol II and pSPT5 occupancy on the MYC promoter using pPol II (S2) or pSPT5 specific antibody in CHiP qPCR analysis (**Figure 5, K,L**), altering MYC transcription at the TSS and downstream gene body towards the CPS (**Figure 5,K**). PTEN loss of function results in massive increase in global levels of nRNA and upon treatment with LCI133 we observe rapid decline, or “nose dive”, in the levels of global nRNA within EAC cells (**Figure 6, E**). Moreover, we observed marked alterations in TS of specific gene sets (RNAseq) in PTEN shRNA KD EAC cells treated with LCI133 including MYC target genes, genes involved in the control of mitosis, and genes controlling RNA processing including splicing (**Figure 6, F-G, S11, A/B**). Phenotypically, PTEN shRNA EAC cells were more sensitive (IC_50_ and apoptotic response) to LCI133 than PTEN wild type EAC cells (**Figure 5, A-B, 6, C-D**).

**Figure 5.**
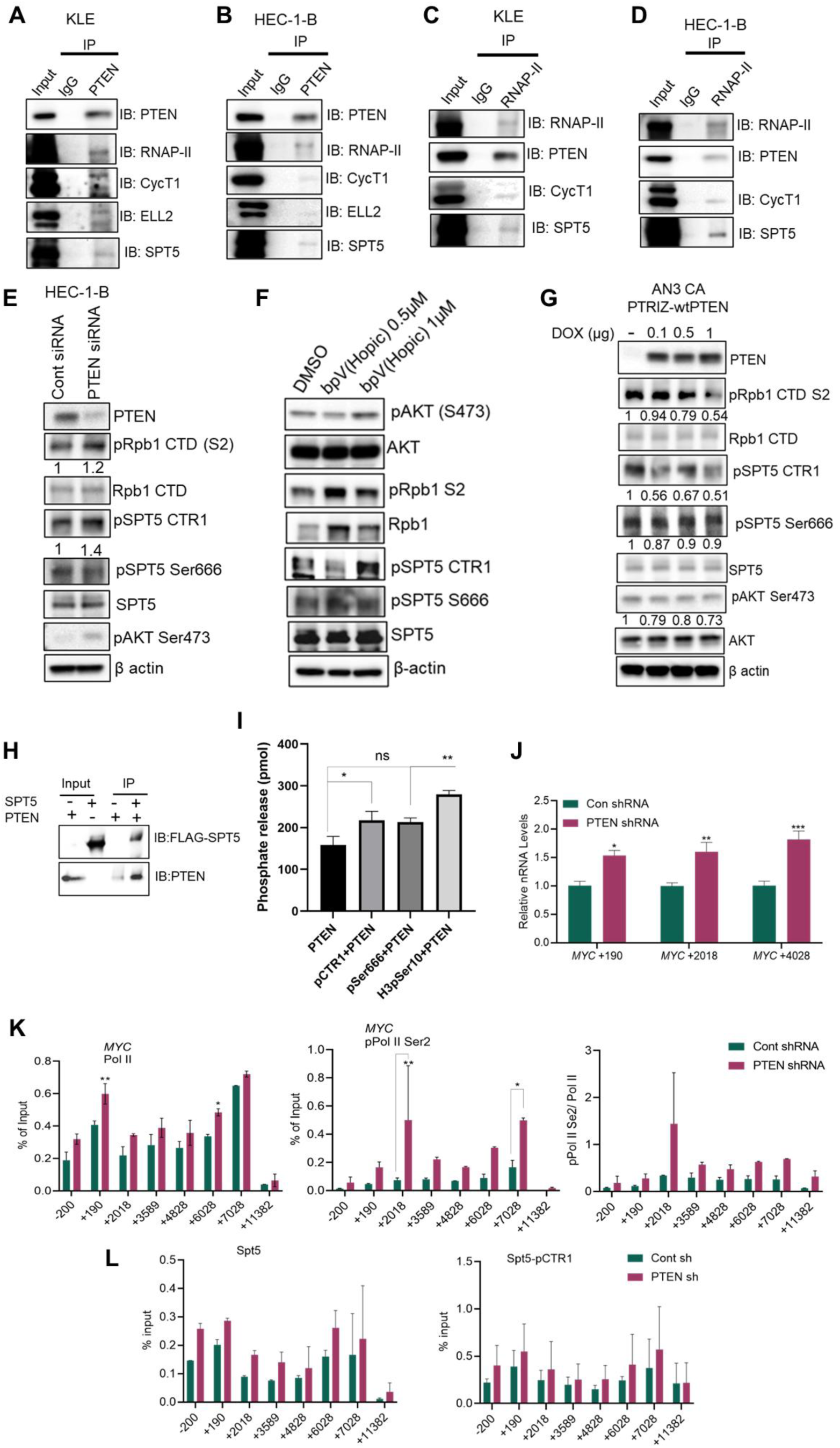
PTEN regulates transcriptional elongation, is bound to SEC proteins, dephosphorylates SPT5, controls pol II/pPol II and SPT5/pSPT5 occupancy on the MYC promoter and regulates global nRNA levels in EAC cells. **A-D**, Co-IP and reciprocal Co-IP of PTEN or RNAP-II followed by Western blot analysis demonstrates a protein-protein interaction between Pol II, PTEN, Cyclin T1, CDK9, ELL2, and SPT5 in PTEN WT KLE (A, C) and HEC-1-B (B, D) EAC cells under resting conditions. **E,** Western blot analysis of PTEN, pRpb CTD, total Rpb1, pSPT5CTR1, pSPT5Ser666, total SPT5, and pAKT Ser473 in control and PTEN siRNA transfected HEC-1-B EAC cells. Numbers below the blot show the quantification of band intensities after normalization to respective β-actin loading control in comparison with siRNA control samples using ImageJ software. **F,** Western blot analysis of pAKT, total AKT, pRpb CTD S2, total Rpb1, pSPT5 CTR1, pSPT5Ser666, and total SPT5 in HEC-1-B cells treated with the indicated concentration of PTEN inhibitor bpV(Hopic) for 24-hours. β-actin served as loading control. **G,** Western blot analysis of PTEN, pRpb CTD S2, total Rpb1, pSPT5 CTR1, pSPT5Ser666, total SPT5, pAKT Ser473, and total AKT in AN3 CA PTRIZ-wtPTEN EAC cells. Numbers below the blot show the quantification of band intensities after normalization to respective β-actin loading control in comparison with no DOX control samples using ImageJ software. **H,** In vitro pulldown assay using immobilized FLAG-SPT5 as the bait and incubated with His-PTEN. The bound protein was subjected to SDS-PAGE followed by Western blotting. **I,** Phosphopeptides derived from Spt5, specifically those containing pSer666 or pCTR1 T806, along with a control histone H3-derived phosphopeptide containing pSer10, were incubated with purified recombinant PTEN or with PTEN alone, as indicated. The measurement of phosphate release was conducted by colorimetric malachite green phosphate assay. Error bars represent the mean ± S.E.M from three biological experiments. Data were analyzed using a two-way ANOVA where P value: * p ≤ 0.05 or ** p<0.01. **J,** qRT-PCR analysis of MYC nascent RNA of in Con shRNA vs PTEN shRNA HEC-1-B EAC cells. The numbers below the graph show the position of primers from the TSS (bp). Error bars represent ± S.E.M from two biological experiments. Data were analyzed using a two-way ANOVA where P value: ** p ≤ 0.01. **K** and **L,** Stable shRNA Con or PTEN knockdown in HEC-1-B EAC cells was assessed by ChIP qPCR analysis for relative occupancy of total Pol II/pPol II **(K)** and total SPT5/pSPT5 CTR1 **(L)** within the gene body of MYC. Error bars represent ± S.E.M from two biological experiments. Data were analyzed using a two-way ANOVA where P value: ** p ≤ 0.01 or * p ≤ 0.05 denotes a significant difference in Con shRNA vs PTEN shRNA.

**Figure 6.**
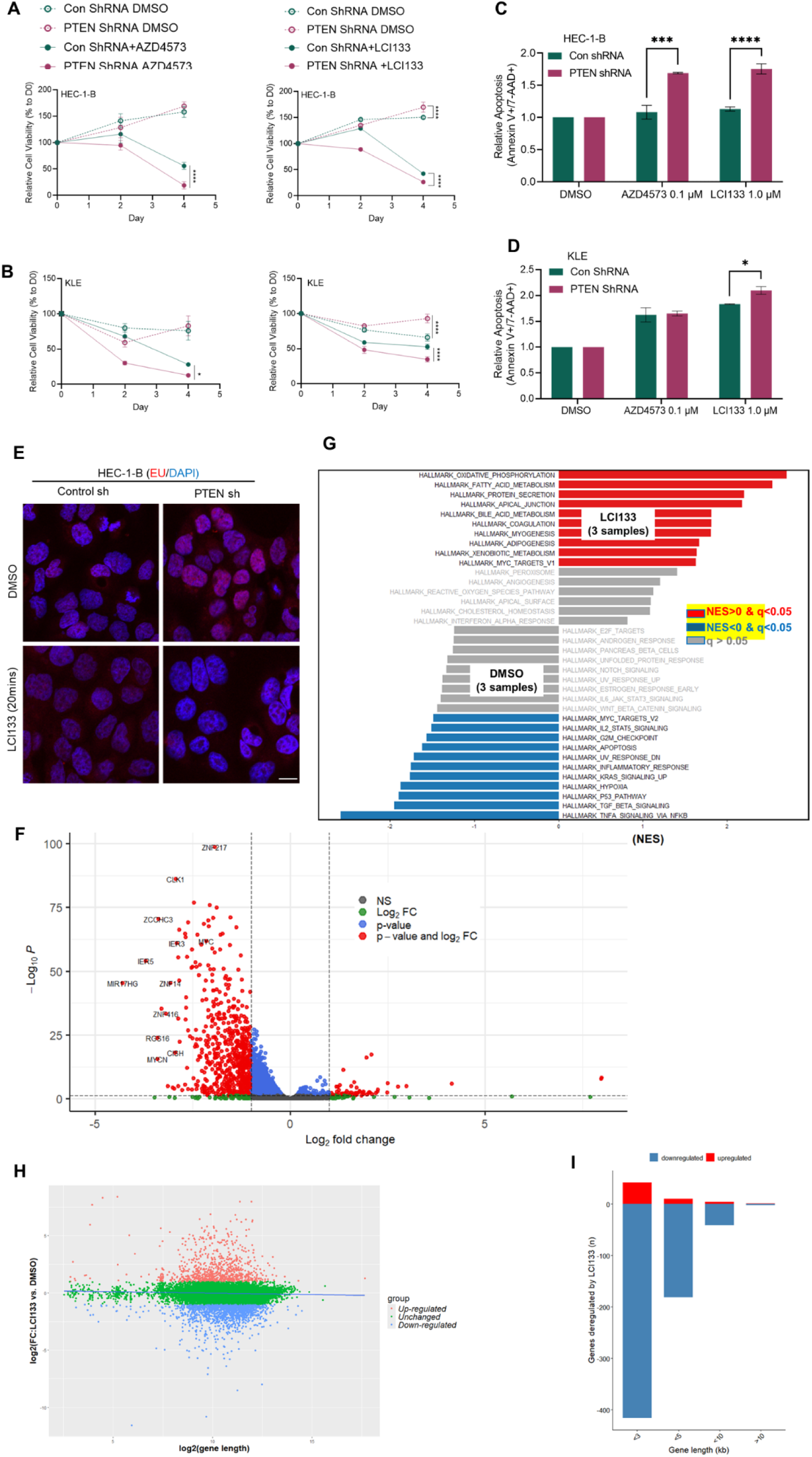
PTEN knockdown or expression in EAC cells affects sensitivity to LCI133 and AZD4573 *in vitro,* controls levels of total cellular nRNA and gene expression. **A** and **B**, The shRNA-mediated knockdown of PTEN in PTEN WT HEC-1-B and KLE cells increases sensitivity to LCI133 and AZD4573. Error bars represent ± S.E.M from triplicate experiments. Data were analyzed using a two-way ANOVA where P value, **** p ≤ 0.0001 or * p ≤ 0.05 denotes a significant difference in DMSO Con shRNA vs PTEN shRNA, LCI133 Con shRNA vs LCI133 PTEN shRNA, AZD4573 Con shRNA vs AZD4573 PTEN shRNA**. C** and **D,** Control and PTEN shRNA stable HEC-1-B or KLE cells treated with DMSO, AZD4573 (0.1 μM), or LCI133 (1.0 μM) and assessed by flow cytometric staining of Annexin V/7-AAD at 48 h. Data represent the mean ± S.E.M. of 3 technical replicates per condition from two independent experiments. **E,** HEC-1B cells expressing PTEN shRNA vs control shRNA were evaluated for levels of total nRNA after LCI133 treatment by confocal microscopy. **F,** HEC-1B PTEN shRNA cells showing differentially expressed genes (DEGs); effect of LCI133 vs DMSO treatment displayed in a volcano plot. The two dashed lines represent the significance thresholds (FDR < 0.05 and fold change > 2). **G,** Gene set enrichment analysis (GSEA). Comparing treatment with LCI133 vs DMSO control in HEC-1B PTEN shRNA transduced EAC. Biological functions and pathways annotated by GSEA hallmark gene signatures are enriched in the LCI133 vs DMSO treatment condition. A negative normalized enrichment score (NES; blue) or positive NES (red) indicates that the gene set is significantly enriched among genes upregulated in the condition of interest, either DMSO or LCI133 treatment. **H,** A scatter plot illustrates the correlation between differential gene expression [log₂(fold change, in HEC-1-B PTEN shRNA treated with LCI133 vs treatment with DMSO)] and gene length [log₂(base pairs)]. Pearson’s correlation analysis yielded a correlation coefficient of −0.04 and a p-value of 2.308 × 10⁻¹⁰. Genes were categorized as upregulated, downregulated, or unchanged based on the following criteria: fold change (FC) > 1, FC < 1, and otherwise, respectively. **I,** Number of genes up- or downregulated relative to gene length. DEGs are divided into four gene-length intervals (x-axis), and the number of genes in each group is shown on the y-axis.

Conversely, the exogenous stable doxycycline-induced expression of PTEN in AN3 CA cells results in a decrease in pSPT5 phosphorylation (**Figure 5, G**) and a restoration in normal highly controlled TS regulatory mechanisms in particular SPT5 CTR1 a component of the SEC and eliminated the build-up of total global nRNA (**Figure S10, E).** Phenotypically, restoring wild type PTEN to PTEN shRNA transduced HEC-1-B cells significantly lowered their sensitivity (IC_50_ and apoptotic response) to treatment with LCI133 or AZD4573 (**Figure S10, B**). Reversal of PTEN loss of function affects EAC Pol II (S2) occupancy beyond the cleavage and polyadenylation site (CPS) of MYC gene bodies indicative of a possible defect in termination. PTEN loss of function results in the augmented rate of transcription and accumulation of Pol II at distal MYC promoter regions including the termination zone (TZ) (+6028-7028) (**Figure 5, K**). From these data, we conclude PTEN reconstitution restores the normal highly controlled TS regulatory mechanisms particularly in the elongation phase controlled by pSPT5 CTR1. We demonstrate that PTEN knockdown in HEC-1-B and KLE cell lines results in: **1**) alterations in MYC transcription at the TSS and downstream gene body towards the CPS (**Figure 5, J**) **2**) increase Pol II/pPol II and SPT5 occupancy on the MYC promoter using pPol II (S2) or pSPT5 specific antibody in CHiP qPCR analysis (**Figure 5, K,L**) **3**) Increased sensitivity to LCI133 and AZD4573 (**Figure 6, C-D**) **4**) dramatic increase in apoptosis in PTEN KD EAC cells to LCI133 and AZD4573 (**Figure 6,C-D**). Herein, we demonstrate for the first time, that wild type PTEN associates with Pol II, cyclin T1/CDK9, ELL2 and SPT5 in HEC1-B and KLE in EAC cells and the suppression of PTEN function using shRNA, mutation or deletion is necessary and sufficient to determine sensitivity to LCI133 and AZD4573 treatment (**Figure 6, A-D**). Using SPT5 phosphorylation site phosphopeptides, epitope tagged mammalian expression constructs of SPT5 CTR1, CTR2 and KOWx-4/5 linker, a PTEN inhibitory chemotype and recombinant PTEN we demonstrate PTEN directly dephosphorylates SPT5-CTR1 but not S666 in the KOWx-4/5 linker region of SPT5 *in vitro* and in cell-based assays and controls levels of nRNA on the MYC promoter in EAC cells (**Figure 5, F,G,I**). As PTEN LOF augmented MYC TS of nRNA in EAC cells, we postulated LOF PTEN disrupts the orderly control of elongation, fueling TA and therefore the sensitivity to LCI133 and other TS inhibitors. Further, we posit aberrant augmented and deregulated TS/TA could increase the content of nascent non-transcribable RNA (nRNA) and the marked rate of global nRNA decay of oncogenic or survival transcripts other than MYC and MCL-1, upon LCI133 or AZD4573 treatment could be causal in the observed augmented apoptotic sensitivity termed “**transcriptional crisis**”. This is supported by our determination of increased levels of apoptosis in PTEN mutant vs wild type EAC cells (**Figure S3,A-D**) and in HEC-1-B or KLE cells treated with PTEN shRNA KD vs control shRNA KD or PTEN null cells reconstituted with wild type PTEN (**Figure 6, C-D**) (**Figure S5, A-F**). Moreover, PTEN associates with SPT5 and recombinant PTEN dephosphorylates SPT5 CTR1 but not KOWx-4/5 (**Figure 5, H/I**).

### PTEN shRNA knockdown in PTEN WT EAC cells increases sensitivity to LCI133 and AZD4573 *in vitro* and controls levels of total cellular nRNA and gene expression

PTEN KD in WT EAC cells resulted in augmented sensitivity to CDK9 inhibition, but not the inhibition of CDK7, 11 and 12 (**Figure 6, A-D; S11**). These results are consistent with our data using the profiling EAC using CRISPR screens for pathway dependency map for CDK9, CDK4 and Aurora kinase where CDK9 shows (Figure S2, B). PTEN loss of function results in massive increase in global levels of nRNA and upon treatment with LCI133 we observe rapid “**nose-dive**” in the levels of global nRNA in EAC cells (**Figure 6, E**). Moreover, we observed marked alterations in TS of specific gene sets (RNAseq) (57 genes upregulated and 640 downregulated) associated with the loss of function of PTEN via the treatment of PTEN shRNA in EAC cells and treated with LCI133 including but not limited to: **1**) MYC target genes **2)** genes involved in the control of mitosis **3)** genes controlling RNA processing including splicing (**Figure 6, F-I S10, A/B**). Of note, it is observed that a predominance of transcripts downregulated are less than 10 kb in size (**Figure 6, H,I).**

### Isogenic PTEN LOF in EAC increases *in vivo* sensitivity to LCI133

We compared the *in vivo* sensitivity of PTEN WT EAC cells to those stably transduced with PTEN shRNA vs control shRNA. We implanted these isogenic EAC cell lines into NSG mice and treated with LCI133 vs vehicle control as described in the methods section (**Figure 7, A-D**). Treatment with LCI133 40 mg/kg IP daily for 21 days resulted in a marked suppression of tumor growth selectively in mice implanted with PTEN shRNA KD EAC cells and not shRNA controls. Moreover, treatment with LCI133 was not associated with toxicity as animal weight remained normal in these mice. We examined the *in vivo* pharmacologic inhibitor of two known CDK9 target genes, MYC and MCL1 and noted an abrogation of both oncogenic proteins on day 21 of treatment. Pathological examination of multiple organs tissues in these mice e.g. liver, kidney, heart and spleen were normal by pathological evaluation.

**Figure 7.**
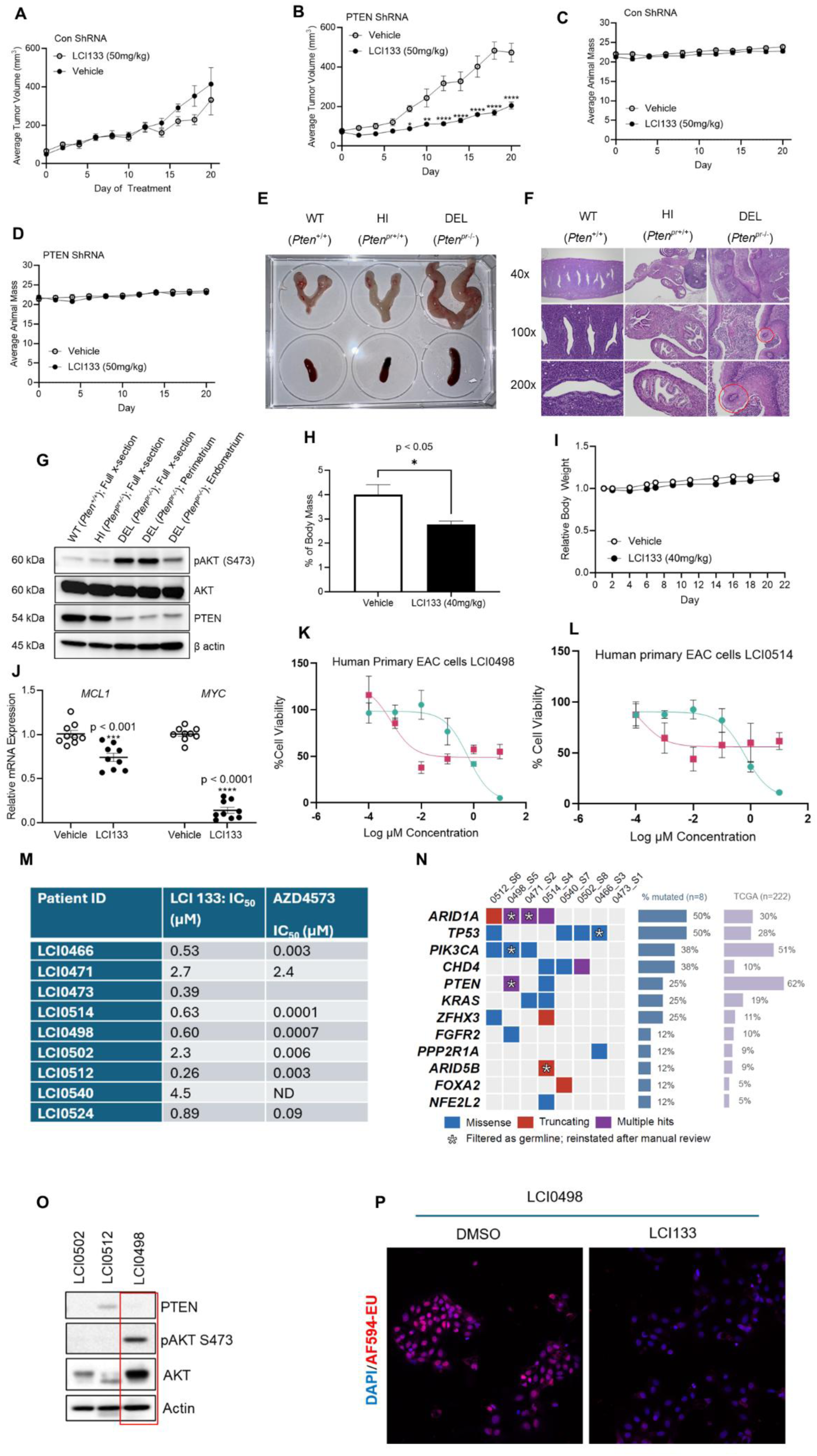
LCI133 is tolerable and efficacious *in vivo* against PTEN shRNA KD EAC cells xenograft and genetically engineered mouse model (GEMM) models of EAC (E-J), in PTEN KD EAC (A-D) or in *ex vivo* primary PTEN mutated EAC patient derived EAC cells (K-P). Beginning 13 days and 17 days after HEC-1-B PTEN shRNA or Con shRNA xenograft implantation, mice from each cohort were randomized into two groups and treated with vehicle (n = 7) or LCI133 (n = 8) for 21 days as described in the methods section. (**A**, **B**) Quantification of tumor volume and (**C**, **D**) animal weights during treatment with LCI133 or vehicle control (50 mg/kg/dose/day x 21 days) or at humane endpoint, when appropriate. Values are mean ± SEM and were analyzed by two-way ANOVA or student’s t test where ****P < 0.0001, ***P < 001, **P < 0.01 (Graph Pad Prism). **E-F**, A conditional PTEN knock out model (PTEN^fl/fl^) and a progesterone receptor promoter driven cre recombinase (PR^cre+^) to induce a knock-out of one or both PTEN alleles in the endometrium. PTEN^fl/fl^ mice were crossed with PR^cre+^ to produce: 1) PTEN^fl/wt^ x PRcre+ (PTEN haploinsufficient; HI) or 2) PTEN^fl/fl^ x PRcre+ (PTEN homozygous deleted; DEL) compared to 3) PRcre+ only (no flox PTEN wildtype; WT) mice. This PTEN^-/-^ DEL mouse recapitulates *in situ* EAC from 4-8 weeks of age. The female reproductive tract (FRT) was excised at 7-8 weeks of age and mounted for gross comparison with paired spleens (E) or mounted, sectioned, and stained with H&E (F). Representative images are shown of n = 3 animals per genotype. **G**, Western blot analysis of full cross-section of the FRT in PTEN WT, HI, and DEL animals and isolated section of perimetrium and endometrium from PTEN DEL mice probed for PTEN, pAKT (S473), and total AKT with β actin as loading control. **H-I**, At 4-6-weeks-old, PTEN DEL animals were randomized into vehicle or LCI133 (40 mg/kg) groups and dosed daily by intraperitoneal injection for 21 days consecutively. At endpoint, relative FRT/total body weight ratios as an indicator of efficacy (**H**) and animal weight as an indicator of toxicity (**I**) comparing vehicle to LCI133 treated animals were assessed (n = 6 per group). **J,** The pharmacodynamic effects of LCI133 vs vehicle control on primary oncogenic transcriptional targets *in vivo* at the end of tumor treatment. Data represent the mean ± S.E.M. of 3 technical replicates per condition from animals per experimental group were analyzed by two-way ANOVA or student’s t test where P value: **** p ≤ 0.0001, ***p ≤ 0.001, and * p ≤ 0.05. **K-L,** Human primary EAC cells freshly isolated from patient EAC tumor tissue LCI04981(**K**) and LCI0514(**L**) were treated *ex vivo* with either increasing concentration of CDK9i, AZD4573 or LCI133 and evaluated 48-hours later for cell viability by CellTiter-Glo assay. Data represents the mean of triplicate experiments ± SD. **M,** Table showing the IC_50_ value of LCI133 and AZD4573 from freshly isolated primary EAC cells. **N**, Oncoplot of WES data showing 22 mutated genes of TCGA UCEC. Twelve carry a call here, from 32 calls across 7 of the 8 tumors. Nine of the 32 were filtered by Mutect2 as germline and reinstated on review; these are marked on the figure. No matched normal was sequenced, so somatic origin is inferred rather than established. **O,** Western blot analysis PTEN, pAKT S473 and total AKT from freshly isolated EAC tumor cells in PTEN LOF (ID: LCI0498) and PTEN WT (ID: LCI0512 and ID: LCI0502). Beta actin used as a loading control. **P,** Representative confocal microscopy images showing effect of LCI133 (0.5 μM) on the abundance of EU-labeled cellular nascent RNA (magenta) in PTEN mutated human primary EAC cells (ID: LCI0498) compared to DMSO control.

### LCI133 is tolerable and displays antitumor activity in PTEN^fl/fl^ x PR^cre+^ EAC GEMM model

To investigate the antitumor effects of LCI133 in an immunocompetent *in vivo* model we used the well-established uterine cre-recombinase specific homozygous PTEN KO GEMM model in which 100% of homozygous KO mice develop EAC between 4-8 weeks of age We confirmed the ratio of the weight of female reproductive track (FRT)/total animal weight (measure of tumor growth) was increased in the PTEN^fl/fl^ x PR^cre+^ (homozygous conditional knock-out) (DEL) relative to the PTEN^fl/fl^ wild type vs PTEN^fl/wt^ x PR^cre+^ (PTEN haploinsufficent) (HI) (**Figure 7, E**). Homozygous loss of both alleles of PTEN in the endometrium resulted in 3-fold increase in FRT/body weight mass ratio and 6-fold increase in total splenocytes/spleen. Histopathology confirms presence of EAC within the enlarged FRT of all PTEN -/- mice including those in the therapeutic study. Western blot demonstrated a loss of expression of PTEN and an increase p-AKT in the endometrium of PTEN^fl/fl^ x PR^cre+^ animals (**Figure 7, G**). Results show Pten^fl/fl^ x PR^cre+^ mice treated with LCI133 (40 mg/kg/dose/daily) for 22 days display a significant inhibition of FRT/animal weight and tumor volume confirmed by histopathological analysis (**Figure 7, H**) (n =6 per group) (p < .05) and a lack of overt toxicity as determined by no loss of body weight or an assessment of an animal toxicity scoring rubric (**Figure 7, I**) and normal histopathology of liver, kidney, lung, heart as reviewed by a pathologist (data not shown). In addition, it was observed, LCI133 pharmacodynamically shuts down the transcription of MCL-1 and MYC mRNA in these EAC tumors *in vivo* (**Figure 7, J**). <u>Importantly</u>, LCI133 does <u>not display any evidence of toxicity in mice</u> treated daily for over 21 days **(Fig. 6, C, D and I).**

### PTEN mutational status in EAC patient derived primary tumor cells is associated with marked sensitivity to LCI133, AZD4573 and LCI133 abrogates nRNA synthesis in PTEN LOF EAC cells

EAC cells isolated from fresh tumors tissues, were assessed for: **1**) Sensitivity to LCI133 or AZD4573 (**Figure 7, K-M**) **2**) Effect of LCI133 on nRNA levels in a PTEN LOF primary EAC patient, LCI0498(**Figure 7,O, P**). We performed whole exome sequencing (WES) to evaluate the PTEN mutational status in EAC isolates except LCI0524 and those subjected to LCI133 treatment. We performed LCI133 IC_50_ determinations and nRNA quantitation. **Fig. 7**.shows the oncoplot with 12 significantly mutated genes including PTEN in a cohort of 8 EAC patients. Consistent with TCGA data, our WES data demonstrate that *PTEN* is co-mutated with its known partner *ARID1A, KRAS* and *PIK3CA* in specific EAC patient samples. Consistent with the WES data, patient LCI0498 EAC cells display loss of mutation of PTEN and abrogation of PTEN protein expression and an increased in pAKT (**Fig, 7 O**). LCI0498 EAC cells displayed high levels of total nRNA and LCI133 treatment *ex vivo* resulted in a rapid near complete abrogation of nRNA synthesis in these cells associated with nM sensitivity to this agent. Specifically, the fluorescence signal from EU labeled nRNA was 80% reduced 20 min after exposure to LCI133 vs the DMSO control indicating rapid and profound suppression of global nRNA transcription paralleling results in bulk RNAseq analysis, as observed in our isogenic models of PTEN LOF vs WT EAC cell lines. LCI133 exhibited nanomolar potency in patient EAC cells isolated *ex vivo* from PTEN mutated EAC cells LCI0514(630 nM) and LCI0498(600 nM) as compared to PTEN WT human EAC cell lines (2.7 µM) (**Figure 7, M).** These data parallel our results in isogenically PTEN shRNA KD EAC cell lines where PTEN LOF is causally linked to nRNA synthesis, augmented global TS (TA), nM sensitivity to LCI133 and CDK9i and marked ablation of nRNA by these chemotypes (**Figure 6/7**).

## Discussion

Treatment of cancer with a single targeted therapeutic agent is invariably associated with the development of resistance followed by relapse ^4,25–29^. Herein, we describe the intentional use of scaffold-merging strategies (no linkers or high-through-put screens) combined with pharmacophore fragment-based computational modeling to engineer MTDLs which will bind selectively to CDK9, CDK4/6 and AURK3 with high affinity, LCI133 and other chemotypes that also bind to PI3K and/or BRD4, LCI132. The rationale for targeting BRD4 and PI3K relates to evidence in the literature that resistance to single targeted therapeutics involves an epigenetic process termed “kinome adaptation” which is blocked by BRD4 inhibition ^30,31^ and apoptotic resistance which is regulated by an important central regulatory kinase involved in apoptotic sensitivity and cell survival, the PI3K/AKT pathways ^32–35^. We used structural datasets and computational chemistry to quantitate target specific binding affinity (free energy scores e.g. (GLID-binding scores) to determine the strengths of molecular interactions between each chemotype and regions of each cognate target molecule binding site e.g. BRD4, H3K27, acetyl lysine; PI3Kα, V851 with morpholino oxygen. etc. in order to engineer one chemotypes to inhibit 3 different molecular targets. To convert LCI132 to LCI133, we neutralized LCI132’s binding of the morpholino oxygen to the PI3Kα, V851 site within the ATP pocket active site ^7^. We computationally altered LCI132 with different “**warheads**” in order to modulate target selectivity and tested them in cancer models as illustrated in this and other manuscripts^1,2^. We posit that each chemotype of a given defined target selectivity could be efficacious against different SL in different malignancies alone or in combination (to be tested). This polypharmacology strategy challenges the failed, long-established “one drug/one target” paradigm ^4,7,25,27,28^.

Despite advances in the molecular characterization of EAC, the rational application of targeted therapy towards common and frequent mutations in this disease, e.g. PTEN mutation, are limited ^10^. PTEN/PI3K/AKT pathway aberrations occur in 80% of endometrial cancer patients. Importantly, the TCGA demonstrated PTEN loss of function mutations are present in both high-grade and low-grade EAC tumors distributed within the POLE, MSI and copy number low subgroups which have significant mortality ^9^. Despite the 80% incidence of PTEN mutations in EAC in both the endometrioid and non-endometrioid high-grade and low-grade variants, PTEN status is not used to prognosticate and/or for the design of therapy for this malignancy^36^.

Mammalian transcription is a complex process involving the highly dynamic formation of multi-protein complexes involving p-TEFb and other proteins at different stages e.g. initiation, pause release, re-initiation, elongation, polyadenylation, termination, maturation of nascent mRNA and splicing ^37^. Pol II and the mediator transcriptional complex include highly defined temporal-spatial phases involving specific kinetic phosphorylation/dephosphorylation dependent changes in multiple components including, PTEN, CDK7, CDK9, PP2A, PP4, RNA Pol II, INTSA, SPT5, PP1, ALL3, ELL2, PNUTS, NELF, PAF1C, RTF1, CDK12/13, etc.^37^. Importantly, CDK9/p-TEFb phosphorylates serine 2 within the C-terminal domain (CTD) of Pol II to promote pause release, transcriptional elongation and global TS of proto-oncogenes such as MYC, MCL1 and other transcripts involved in carcinogenesis and cell growth ^21,22,38^. It is reported that PTEN interacts with the Pol II transcriptional machinery via its C2 domain and via its interaction with chromatin modulates gene transcription by the genome-wide redistribution of Pol II occupancy on promoter enhancer elements ^12^. This regulation of gene expression is dependent upon PTEN’s catalytic phosphatase activity ^37^. PTEN is thought to impact Pol II pause duration and release and the elongation rate thereby affecting global transcription ^12^. PTEN binds to promoter elements in chromatin and negatively regulates genes involved in transcription e.g. AAF and POL2RA to name a few ^13^. Our results indicate loss of PTEN results in an increase in SPT5 phosphorylation of a stretch of phosphorylated serine/threonine residues within the CTD of SPT5, CTR1 domain (T806) but not on S666 in the KOWx-4/5 linker domain. We confirmed the direct dephosphorylation of pCTR1 by PTEN using recombinant PTEN. The pSPT5 CTR1 dephosphorylation has been previously reported to be carried out by PP1 and it is reported to regulate the orderly process of elongation and transcriptional termination ^39–41^. Our data provide direct and indirect evidence for PTEN’s regulation of the fine-tuned elongation phase of TS via the potential control pSPT5 CTR1.

Using PTEN isogenically manipulated EAC cells we demonstrate that PTEN loss of function is *necessary and sufficient* to result in a dramatic increase in total nRNA TS compared to PTEN wild type EAC cells. Moreover, inducible PTEN reconstitution of PTEN null EAC cells rapidly drops levels of nRNA TS. Our data reveal, a massive augmented decay in the global levels of nRNA in PTEN shRNA KD cells upon exposure to LCI133, far above that observed in isogenic EAC cells with WT PTEN. This is associated with a marked alteration in gene expression in particular gene sets involving the regulation of RNA processing and splicing and the induction of marked apoptosis.

From these combined data, we conclude the abrogation of MYC and MCL1 gene expression is neither *necessary nor sufficient* to encode the apoptotic sensitivity to LCI133 or CDK9i in PTEN loss of function isogenic EAC models suggesting other suppressed genes within the 525 observed in our RNAseq dataset may account for this potential SL. Other observations from our isogenic models support a **causal role for PTEN loss** of function in the sensitivity to LCI133 including these phenotypes: **1**) a decrease in IC_50_ for AZD and LCI133 in PTEN mutant or PTEN shRNA isogenically manipulated EAC cell lines **2**) Increase in the apoptotic response to LCI133 and AZD4573 in wild type EACs treated with PTEN shRNA KD **3**) EAC cells with catalytic dead mutant PTEN (G120R) exhibit augmented sensitivity and apoptosis to LCI133 and AZD treatment suggesting loss of PTEN catalytic phosphatase activity is necessary and sufficient to drive sensitivity to LCI133 and AZD4573 **4**) Co-immunoprecipitation of PTEN with Pol II, cyclin T1, ELL2 and SPT5, part of the SEC in EACs **5**) PTEN loss results in an increase in Pol II occupancy on the distal gene body of the MYC promoter including the termination zone **6**) PTEN associates with and is required for the dephosphorylation of pSPT5, a necessary event in the highly tuned control of orderly elongation and termination phase of the TS cycle **7**) Suppression of PTEN via shRNA results in increased Pol II/serine 2 phosphorylated Pol II occupancy MYC promoter region on the distal gene body beyond the cleavage and polyadenylation site (**CPS**) suggesting PTEN may function at the elongation and/or termination stage of TS **8**) reversal of PTEN loss of function in EAC results in a decrease in Pol II (S2) occupancy beyond the cleavage and CPS site of MYC gene bodies indicative of a defect in termination **9**) knockdown of PTEN in PTEN WT EAC cells results in the control of ELL2 expression a component of the TS SEC **10**) Isogenic LOF PTEN results in augmented TS consistent with TA and treatment of PTEN LOF EAC cells induces a rapid and dramatic drop in global nRNA transcripts compared to PTEN WT EAC cells and augments apoptosis. **11**) Reconstitution of WT PTEN in PTEN null EAC cells results in suppression of TS and TA and loss of sensitivity to LCI133 **12**) The rapid and total abrogation of total nRNA and nM sensitivity to LCI133 was recapitulated in *ex vivo* propagated PTEN mutated primary patient derived EAC cells. These data support a model implicating PTEN loss of function in the control of global nRNA TS (TA) and document LCI133 induces a massive drop in total nRNA in particular low molecular weight (< 10kb) transcripts involved in RNA splicing a phenotype which encodes a marked apoptotic sensitivity to LCI133 in EAC cells. These results suggest PTEN LOF is a mechanistic biomarker for sensitivity to LCI133 and/or other CDK9 inhibitors in EAC cancer cells.

In summary, our findings suggest that LCI133 is a promising lead chemotype for development as a polypharmaceutical targeted therapy for PTEN loss of function EAC. As outlined above, we provide experimental evidence for a novel mechanism by which PTEN loss of function in EAC cells defines sensitivity to LCI133 and other CDK9 inhibitors. Finally, treatment with LCI133 daily for 21 days shows efficacy and not evidence to toxicity.

## Limitations of study

Although *in vitro* and *in vivo* models show promising results for the potency and safety of LCI133 in PTEN loss of function EAC cells, these models cannot fully reproduce the complexity of human cancer physiology. It’s clinical safety and efficacy in humans with PTEN LOF EAC or other PTEN LOF cancers will require further study and clinical trials.

## Author contributions

D.D., K.M. and D.P conceived the hypothesis and wrote the manuscript. D.D, C.M, D.P, K.M. D.R, V.M, contributed to experimental design and data analysis. D.D. provided financial support. R.D., V.M., Q.Z., D.P., C.M. and H.D. performed the *in vitro/in vivo* experiments. K.M. and D.D. performed target selection and *in silico* design and B.Y. provided synthetic organic synthesis. J. L. and D.F. reviewed the manuscript. W.N., A.P., E.C., J.B. provided an analysis of EAC clinical relevance and provision of EAC tumors from OR. W.S. P.Y and K.D. for biostatistical or bioinformatics support, respectively.

## Acknowledgements

The authors cite grant support from the NIH/NCI RO1 funding to D.L.D., CA215651, CA172513 and the Wake Forest Baptist Cancer Center support grant P30CA012197 (R.M.), Ovarian cancer grant, NIH/NCI Stepping Stones (SS) program (NIH/DTP 237164). D.L.D is supported by Leukemia and Lymphoma Foundation TR grant and start-up funds from LCI/WFBCCC and the generous support of The Leon Levine Foundation. ACS IRG and K12 for K.M. We thank Dr. Caudell for pathology support. We thank Dr. Robert Fisher for providing antibody specific for the phosphorylated pSPT5 CTR1 and KOW linker regions (S806 and S666) reactive to a stretch of S/T phosphorylated residues in this domain and critical discussions on this project. We acknowledge the support from the AtriumHealth/Levine Cancer Center/Wake Forest Cancer Center core facilities including the Biorepository, Flow Cytometry, Dru Discovery and Genomics cores (WFSM/CCSG;P30 CA01209) for the support on this project.

## Materials and Methods

### Chemical biological *in silico* design, synthesis and testing of multitargeted chemotypes, LCI132, LCI133 and LCI136

Herein, we used chemical biological approach and *in silico* methods to design small molecules to selectively inhibit specific targets CDK4/6/9 and Aurora kinase plus or minus PI3K or BRD4. LCI132 binds to PI3K-BRD4-CDK4/6/9 and AURK, LCI133 to CDK4/6/9 and AURK but not PI3K or BRD4 and LCI136 to PI3K-CDK4/6/9 and AURK but not BRD4 ^7^. These multitarget inhibitors were synthesized using standard carbon-carbon (C-C) and carbon-nitrogen (C-N) coupling reaction chemistry and 2-morpholino benzopyranone (BP) scaffold and the active pharmacophore moiety 2-aminopyrimidine^7^. An analysis of the crystal structure of the Ribociclib-CDK6 and alisertib-AURKA complexes show that the 2-aminopyrimide moiety is critical because it binds to hinge regions of the CDK6 and AURK A/B active sites. X-ray crystal structural analysis revealed that amine (NH) and nitrogen three bonds connectivity played a bidentate interaction with hinge residues (2-amidopyridine for CDK9 (Cys-106), the 2-aminopyrimidine for CDK6 (Val101) and for AURKA (Met695). In silico modeling of triple inhibitors was based on the binding patterns made with FDA approved CDK4/6i, Ribociclib; AURKAi, alisertib and a CDK9i, AZD4573, and other 2-aminopyrimide analogs (11). LCI132 and LCI133 are positional isomers of LCI133 (MW: 578.67 Da), while LCI136 (MW: 579.66 Da) is a bioisostere of LCI133, which possesses a pyridine ring system between BP and pyrrolopyrimide scaffolds (Figure 1)^5^. Of importance, we chemically engineered LCI132 to bind to BRD4, based on an analysis of x-ray crystal structures, the LCI132 phenyl ring must be in the para 8 position relative to the phenyl moeity in order to bind within the hydrophobic pocket of BRD4 ^1^. In contrast, LCI136 and LCI133 differ in that the LCI136 was designed to bind to PI3K via the introduction of a warhead that which forms a strong hydrogen bond between the morpholino oxygen of LCI136 and the canonical V851^1^. The IC_50_ measurements for inhibition of BRD4 were performed using Alpha Screen assays from Reaction Biology (Hershey PA) on a set of His-tagged bromodomains against a tetra-acetylatedhistone H4 peptide (H4K5ac/8ac/12ac/16ac-Biotin) ligand. PI3K isoforms, CDK4/6, CDK9 and AURK A/B kinase activities were measured using a γ-P32-ATP-derived kinase assay from Reaction Biology ^7^. Kinome selectivity profile for LCI133 against 468 kinase shows inhibition of 4.7% of all kinases as reported^4,5,7^.

### Tissue culture, cell lines, and reagents

The human EAC cell lines HEC-1-A, HEC-1-B, KLE, (PTEN wild type) and AN3 CA, RL95-2, Ishikawa, HCI-EC-23 (PTEN null or mutated) and the human bone marrow stromal (HS5) and embryonic kidney (HEK293T) cells were obtained from ATCC (Manassas, VA)(9), Ishikawa was obtained from Millipore-Sigma (Burlington, MA) and HCI-EC-23 was obtained from Applied Biological Materials Inc. (Richmond, BC, CA). All cell lines were authenticated by short tandem repeat DNA profiling at the respective cell bank or by IDEXX BioAnalytics (Columbia, MO) and maintained as recommended by the manufacturers. AN3 CA, HEC-1-B, and Ishikawa were cultured in EMEM 30-2003 (ATCC) with 10% FBS (Thermo Fisher) and 1% penicillin–streptomycin (P-S; Thermo Fisher). RL95-2 was cultured in DMEM: F12 30-2006 (ATCC) with 10% FBS, 1% P-S, and 0.005 mg/ml insulin. KLE was cultured in DMEM: F12 (ATCC) with 10% FBS and 1% P-S. HEC-1-A was cultured in McCoy’s 5A Medium 30-2007 (ATCC) with 10% FBS. HCI-EC-23 was cultured in RPMI-1640 30-2201 (ATCC) with 10% FBS and 1% P/S according to Rush et al (12). 293T was cultured in DMEM (ATCC) with 10% FBS and 1% P/S. All complete media formulations were stored at 4°C, and cell cultures were maintained in standard cell culture conditions at 37 °C with 5% CO2.

### Cellular thermal shift assay (CETSA)

A cell-based thermal shift assay was performed as described (13) using the RL95-2 EAC cell line as a measure of ligand, LCI133 physical binding to CDK9, CDK4/6 and AURK A/B kinases. EAC RL-95 cells were seeded at 10 x 10^6^ cells in a 10cm plate. Cells were treated with either 0.5 uM of LCI133 or 0.1% DMSO for 24 h. After treatment, cells were pelleted by centrifugation at 400 x g for 5 minutes followed by two-time wash with PBS. Cells were then resuspended in PBS+PI buffer 50 ul resuspended and aliquoted into 7 tubes. Then 6 aliquots for each condition were subjected to heating on a thermocycler set to a specific temperature ranging from 37-62^0^C with a shift of 50°C steps between samples for 3 minutes followed by cooling to room temperature for 3 minutes. Cells were snap frozen in liquid nitrogen and subsequently thawed and this process was repeated three times. Samples were resolved on 4-12% gradient Novex Tris-Glycine acrylamide gels (Invitrogen) by loading an equal volume of lysates for each condition. The gel was transferred to a nitrocellulose membrane (ThermoFisher) and blots were performed to assess the level of proteins stabilized by each specific chemotype. CETSA results were confirmed with Nano-BRET analysis for LCI133 and each molecular target.

### IC_50_ cell viability assay

EAC or normal non-transformed bone marrow stromal (HS5) cells were plated in 96-well plates at a density of 0.5-2 x 10^4^ cells/well in 100 µl of the respective media and incubated overnight to attach. Cells were treated with DMSO (0.1% final) or increasing concentrations of single agent inhibitor control or LCI132, LCI133, or LCI136 and incubated at 37 °C for 48 hours. 100 µl of CellTiter-Glo luminescent cell viability reagent (Promega, Madison, WI) was added to each well and the cells were incubated for 10 minutes at room temperature (RT). Luminescence was detected with a CLARIOstar Plus microplate reader (BMG Labtech, Cary, NC, USA). Percentage of cell viability was calculated from the control (DMSO) and analyzed by nonlinear regression then plotted for dose response using GraphPad Prism (GraphPad Software, Inc., San Diego, CA). Single agents Buparlisib (BKM120), PLX51107, Ribociclib (LEE011), and AZD4573 were purchased from Selleck Chemicals (Houston, TX, USA). HEC-1-B control shRNA or PTEN shRNA cells were plated at 10000 cells per well in a 96 well plate and grown in 3 replicates per condition then exposed to CDK9, CDK7, CDK12 or CDK12 inhibitors on the following day. At indicated times, 100 µl of CellTiter-Glo luminescent cell viability reagent (Promega, Madison, WI) was added to each well and the cells were incubated for 10 minutes at room temperature (RT). Luminescence was detected with a CLARIOstar Plus microplate reader (BMG Labtech, Cary, NC, USA). Percentage of cell viability was calculated by normalizing from day zero and then plotted the viability plot using GraphPad Prism (GraphPad Software, Inc., San Diego, CA).

### Colony formation assay

Due to differences in cell size and proliferation rate, AN3 CA and HEC-1-A cells were seeded at 2 x 10^3^ and 10 x 10^3^ cells per well, respectively, and treated with one-time dose of DMSO (0.1% final) or the indicated concentrations of single agent inhibitor control or LCI132, LCI133, or LCI136 and incubated for 2-3 weeks under standard conditions. Following incubation, media and floating cells were aspirated and discarded, wells were washed twice with PBS, samples fixed with MeOH for 10 minutes in the dark and stained with Crystal violet for 15 minutes at room temperature. Plates were washed twice in filtered water, left to dry overnight, and imaged with a Nikon D3500 single lens camera. Experiments were performed in duplicate.

### Apoptosis assay

EAC cells were seeded in a 6-well plate at a density of 8 x 10^5^ cells/well in 2 mL of the respective medium and incubated overnight to adhere. Cells were treated with DMSO; (0.1% final), Staurosporine (+; 0.1 µM), AZD4573 (0.1 µM), or the indicated concentrations of LCI132, LCI133, or LCI136 and incubated at 37°C for 24 hours. Cells were collected using TrypLE™ Express enzyme (ThermoFisher), washed and resuspended in Annexin V Binding Buffer (BD Biosciences, Franklin Lakes, NJ), and stained with 2.5-5 µL FITC Annexin V and 5 µL 7-AAD Viability Staining Solution (BD Biosciences) for 15 minutes at room temperature protected from light. Following incubation, the sample volume was raised with additional binding buffer and flow cytometric analysis performed by the Atrium Health Levine Cancer Institute Immune Monitoring Core Laboratory using a LSR Fortessa (BD Biosciences) and FlowJo Software v10.9 for batch analysis (BD Biosciences).

### Cell cycle analysis

EAC cells were seeded in a 6-well plate at a density of 8 x 10^5^ cells/well in 2 mL of the respective medium and incubated overnight to adhere. Cells were treated with DMSO (-; 0.1% final), Ribociclib (+; 0.5 µM), or the indicated concentrations of LCI132, LCI133, or LCI136 and incubated at 37°C for 24 hours. Following incubation in Vybrant Dye Cycle (Invitrogen, Eugeen, OR), samples were immediately subjected to flow cytometric analysis by the Atrium Health Levine Cancer Institute Immune Monitoring Core Laboratory using a LSR Fortessa (BD Biosciences). Univariate cell cycle analysis using Dean-Jett-Fox model was performed in FlowJo Software v10.9 (BD Biosciences).

### RNA extraction, cDNA synthesis and qRT-PCR

EAC cells were seeded in a 6-well plate at a density of 0.5-2 x 10^6^ cells/well in 2 ml media and incubated overnight to adhere. Cells were treated with DMSO (0.1% final), AZD4573 (0.1µM), or LCI133 (1.0 µM) and incubated at 37°C for 2 hours. Following incubation, total RNA was extracted using the RNAeasy Mini Kit (Qiagen, Germantown, MD, USA) according to manufacturer instructions. 1 ug total RNA was reverse transcribed using iscript cDNA Synthesis Kit (Bio-Rad, Hercules, CA, USA) according to manufacturer’s instructions. Amplification of cDNA was performed with PowerTrack SYBR green master mix (Applied Biosystems, Thermo Fisher) on QuantStudio 3 Real-Time PCR System (Applied Biosystems, Thermo Fisher). cDNAs were amplified using custom generated specific primers for MYC, MCL1, RPL30 (Figure S9) Gene expression was normalized to housekeeper RPL30 and calibrated to vehicle controls using 2-ΔΔCt method. nRNA quantitation using 5-ethynyluridine (EU) labeling and immunofluorescence

The Click-iT RNA Imaging Assay (C10330, Thermo Fisher) was used to detect newly synthesized nascent RNA. EU was incorporated into newly synthesized RNA and detected by Alexa Fluor 594 dye according to the manufacturer’s protocol (Thermofisher). Briefly, AN3 CA or stable HEC-1-B PTEN ShRNA KD cells were seeded in complete medium onto glass coverslips in a 6-well plate and incubated to adhere. Cells were treated with DMSO, LCI133 (1.0 µM), or AZD4573 (0.1 µM) then labeled with 1 mM EU for 1 hour. Next, cells were fixed with 3.7% formaldehyde, permeabilized with 0.5% triton-100, and incubated with Click-iT Reaction Cocktail for 30 minutes. The cells were washed once with Click-iT reaction buffer and nuclei were stained by using DAPI (H-1200-10, Vector Laboratories). Fluorescence was visualized with an Olympus confocal microscope (model OX80). where red fluorescence indicates newly synthesized RNA and blue fluorescence indicates DNA/nuclei.

### Nascent RNA capture, quantitation of total nRNA by flow cytometry, reverse transcription of specific transcripts by qPCR and RNAseq analysis

The Click-iT Nascent RNA Capture Kit, for gene expression analysis (C10365, Thermo Fisher) was used to isolate and detect newly synthesized nascent RNA. Untreated or treated AN3 CA or stable HEC-1-B PTEN ShRNA KD cells were labeled or not labeled with 0.5 mM EU for 60 min before collection. 5-ethynyluridine (EU)-labeled nascent RNA was purified according to manufacturer’s protocol (Click-iT Nascent RNA Capture Kit, for gene expression analysis, ThermoFisher, C10365). RNA was quantified using a DS-11 FX+ Spectrophotometer/Fluorometer (DeNovix) and cDNA was synthesized using SuperScript VILO cDNA Synthesis Kit (11756050, Thermo Fisher). Real-time qPCR was carried out with PowerTrack SYBR green master mix (Applied Biosystems, Thermo Fisher) and custom oligonucleotide primers (IDT) on a QuantStudio 3 Real-Time PCR System (Applied Biosystems, Thermo Fisher). The relative level of gene expression was determined using the 2(-DDCt) method and calibrated to the indicated housekeeper gene. The forward and reverse primer pairs for detection by qPCR are shown in the supplemental methods. Levels of MYC specific nRNA were quantitated following EU capture by RT PCR using primers which represent different regions of the MYC promoter +190, +2018 and +4028 distal to the TSS containing intronic and exonic nRNA sequences. For total global nRNA, we used flow cytometric analysis and confocal IF microscopy to quantitate EU-labeled nRNA fluorochrome.

### Western blot analysis

For Western blots, EAC cells were seeded in a 6-well plate at a density of 0.5-2 x 10^6^ cells/well in 2 ml media and incubated overnight to adhere. Cells were treated with DMSO (0.1% final), AZD4573 (0.1 µM), Ribociclib (1.0 µM), or LCI133 (0.5-1.5 µM) and incubated at 37° C for 6 hours. Whole cell lysates were prepared using Pierce RIPA buffer supplemented with Protease/Phosphatase Inhibitor Cocktail from Cell Signaling Technology (CST), Danvers, MA, USA). Protein concentration in cell lysates was determined using Pierce bicinchoninic acid (BCA) assay kit (Thermo Fisher) and absorbance measured using CLARIOstar Plus microplate reader (BMG Labtech). Equivalent clarified lysates were resolved on 4-12% SDS-PAGE, transferred to nitrocellulose membrane(s), and probed with one or more of the following antibodies from Cell Signaling Technology: c-Myc (9402S), Rpb1 (2629S), pRpb1 Ser 2 (13499S), pRpb1 Ser 5 (13523S), Rb (9313), pRb S807/811 (8516), cleaved PARP (9541S), Mcl-1 (39224S), and β-actin (CST 4967S) from Santa Cruz Biotechnology, Dallas, TX, USA # c-47778). Secondary antibodies were chosen according to the species of origin of the primary antibody. Protein bands were detected using the Pierce ECL Western blotting substrate (Thermo Fisher) or Millipore Immobilon Forte Western HRP substrate (EMD Millipore Corporation, Burlington, MA), then imaged using VisionWorks v8.21 software (Analytik Jena US LLC, Upland, CA) on a UVP ChemoStudio PLUS (Analytik Jena). β-actin was used as a loading control for all immunoblots.

### Transcriptional profiling, chromatin immunoprecipitation (ChIP) qPCR and analysis of Pol II and pPol II bound to chromatin upon PTEN LOF

AN3 CA, RL95-2 or stable HEC-1-B PTEN ShRNA KD cells were treated with DMSO, AZD4573 (0.1 µM), or LCI133 (1.0 µM) for 2 hours and were then harvested and processed using SimpleChIP Enzymatic Chromatin IP Kit according to the manufacturer instructions (Cell Signaling Technology, Danvers, MA). In brief, the cells were fixed with 1.1% formaldehyde, quenched by 10% glycine, and lysed in diluted protease inhibitor cocktail. The lysates were enzymologically digested and sonicated in order to shear DNA to form DNA fragments with optimal size of 150-900 bp. 5-8 ug of sheared chromatin was immunoprecipitated with total Rpb1 antibody (CST), pRpb1 CTD Ser2 antibody, IgG antihistone as positive control or no antibody as negative control overnight with rotation at 4 degrees C. A portion of the diluted chromatin was set aside for the INPUT. Diluted chromatin was incubated with anti-Pol II/Rpb1 (2629S), anti-histone H3 (4 µg, Abcam) as positive control, or no antibody as negative control overnight with rotation at 4°C. The antibody binding beads were added and washed according to the manufacturer’s instructions. The samples were treated with DNA-purifying slurry and Proteinase K to purify DNA. Samples were subjected to qPCR as described above using the MYC and MCL1 promoter primers targeting the indicated bases ± transcriptional start site (TSS) and downstream components of these genes including the gene body and termination zones. The forward and reverse primer pairs for detection by RT qPCR and ChiP qPCR are shown in the supplemental methods.

### Subcellular protein fractionation and quantitation of chromatin associated components of TS

Subcellular protein fractionation kit for cultured cells (Thermo Scientific #78840) was employed to isolate cytoplasmic, membrane, nuclear soluble, chromatin bound and cytoskeletal protein extract from HEC-1-B cells stably transduced with control shRNA or PTEN shRNA according to the manufacture’s protocol. In brief, 2 x 10^6^ HEC-1-B were resuspended in 100 µL of CEB buffer and incubated at 4° C for 10 mins with frequent mixing followed by centrifuged at 500 x g for 5 mins. Supernatant was collected as the cytoplasmic fraction and the pellet was resuspended in with 100 µL of MEB buffer, incubated at 4° C for 10 mins followed by centrifugation at 500 x g for 5 min. The supernatant represents the membrane fraction. Next, 50 µL of NEB buffer was added to the pellet and incubated at 4°C for 30 mins, centrifuged 500 x g for 5 mins and the supernatant was collected as the soluble nuclear extract (SNE). The chromatin bound nuclear extract (CBNE) was prepared by adding 5 µL of 100 mM CaCl_2_ and 3 µL of Micrococcal Nuclease to the pellet, vortexed and incubated at 37 °C for 5 mins and again vortexed for 15 mins at highest setting. The mixture was subsequently centrifuged at 16,000 x g for 5 mins and supernatant was collected as chromatin-bound nuclear extract.

### Co-immunoprecipitation

For co-immunoprecipitation, HEC-1B, KLE and dox-inducible PTEN reconstituted AN3 CA cells were grown to 70% confluence, collected and lysed in BC200 buffer (25 mM Tris pH 7.5, 200 mM NaCl, 1 mM EDTA, 0.2% Triton X-100. 0.2% glycerol). Lysates were briefly sonicated and centrifuged to collect lysates. Protein concentration was determined by BCA assay. Immunoprecipitation was carried out using the Pierce Co-Immunoprecipitation kit (Cat #26149). In brief, 1 mg of total protein lysate was incubated with anti-PTEN mAb (6H2.1, Millipore), anti-Rpb1 (CST) or anti-mouse IgG covalently crosslinked to AminoLink Plus resin for 2 hours according to manufacturer’s protocal. Beads were washed three times with lysis buffer and eluted with low pH buffer. The eluted fraction was denatured in 1X sample buffer followed by heating for 5 min. at 95 degrees C. Samples were resolved on 4-12 SDS PAGE, transferred to nitrocellulose membranes and probed with one or more specific antibodies. Protein bands were detected using the Pierce ECL Western blotting substrate or Forte Western HRP substrate Millipore Immobilon (EMD Millipore Corporation, Upland, CA) and imaged using VisionWorks v8.21 software on UVP ChemoStudio PLUS.

### *In vitro* pull-down assays

For *in vitro* pull-down assays, an anti-FLAG IgG antibody or nonspecific IgG was conjugated to magnetic beads and used in *an in vitro* pull-down assay. Recombinant FLAG-SPT5 protein (Origene) were incubated with HIS-tagged recombinant PTEN (ThermoFisher) for a duration of two hours at 4°C followed by the addition of anti-Flag conjugated magnetic beads or control IgG conjugated beads for one hour. Control magnetic beads conjugated to IgG served as a control for SPT5 specificity of binding in the pull-down (lane 4 of Western blot. Subsequently, the magnetic beads were subjected to washing, and the SPT5-associated proteins were eluted and resolved by SDS-PAGE by Western blot analysis.

### *In vitro* phosphatase assays

Phosphatase reactions were conducted in 96-well microtiter plates, utilizing a total volume of 25 µL for each reaction. Each reaction mixture comprised 1.0 mg of purified recombinant HIS tagged PTEN and 100 µM of each phosphopeptides, prepared in a reaction buffer containing 20 mM Tris-HCl (pH 7.5) and 10 mM DTT. The reactions were allowed to proceed for a duration of 60 minutes at room temperature and were subsequently terminated through the addition of 100 µL of malachite green assay reagent (12776, Cell Signaling Technology). Following a 15-minute incubation period at room temperature, the absorbance was measured at 630 nm using a CLAIRIstar plus microtiter plate reader (BMG Labtech). Additionally, a standard curve based on known quantities of inorganic phosphate was generated to facilitate the determination of picomoles of phosphate produced in each reaction.

### Lentiviral production, transfection, and stable shPTEN knockdown (KD) of PTEN expression

Lentiviral production and packaging were performed according to manufacturer’s instructions (Addgene, Watertown, MA, USA). Briefly, HEK293T cells were grown to 75% confluency on a 10 cm tissue culture plate and transfected with a mixture of pCMV-dR8.2 dvpr (Addgene, #8455), VSV.G (Addgene, #14888), and pLKO-PTEN-shRNA-1320 (Addgene, #25638) or scramble shRNA (Addgene, #1864), and lipofectamine 2000 (ThemoFisher; #11668-027) in Opti-MEM and incubated under standard conditions overnight. Media was refreshed and cells incubated again under standard conditions. Virus was harvested at 48 and 72 hours post transfection in individual harvests or a combined harvest where all the individual harvests were pooled. Viral supernatant was centrifuged at 500 rpms for 5 minutes to pellet any packaging cells that were collected during harvest. Viral concentration of the collected supernatant was then measured using Lenti-XTM GoStixTM Plus (TaKaRa, #631280). Supernatant was concentrated using Lenti-X Concentrator (TaKaRa, #631232) then aliquoted and snap-frozen in liquid nitrogen and stored at −80 °C. For PTEN knockdown in HEC-1-B cells, 2 x 105 cells were plated in 2 mL/well of a 6-well plate and allowed to attach overnight. The next morning, media was changed and supplemented with 4 μg/mL polybrene plus 100 mL PTEN or scrambled shRNA lentiviral particle solution and incubated overnight under standard conditions. Medium was refreshed after 72 hours of transduction. Cells were selected with 3 mg/mL puromycin for an additional 72 hours. Cells were grown under standard conditions with puromycin-containing medium refreshed as needed and stable transfection confirmed by immunoblotting.

### Generation of doxycycline (dox) inducible wild type (WT) PTEN in PTEN null EAC cell lines

To generate inducible WT PTEN in EAC cell lines, lentiviral supernatants were produced as mentioned above for pTRIPZ-Wt-PTEN plasmid Addgene (Cat # 206982). PTEN null AN3 CA cells were transduced with pTRIPZ lentiviral supernatant and polybrene (4 μg/ml) for 3 days followed by selecting with 1 μg/ml puromycin for one week. PTEN expression was induced by treating cells with doxycycline monohydrate (Thermo scientific, Cat # J63805-06). For siRNA transfection and transient PTEN knockdown (KD) in EAC cells, 8 x 10^5^ HEC-1-A, HEC-1-B or KLE cells were plated in 2 mL/well of complete culture medium and allowed to adhere overnight. The following morning, culture medium was discarded, wells washed twice, and medium replaced with 0.9 mL/well of phenol-free Opti-MEM (ThermoFisher). 0.1 mL/well Lipofectamine RNAiMax (ThermoFisher; 13778150) master mix containing control shRNA-A (Santa Cruz; sc-37007) or human PTEN shRNA (Santa Cruz; sc-29459) was added dropwise to the appropriate well and incubated for 6 hours under standard conditions. After 6-hour incubation, 1 mL/well of McCoy’s 5A + 20% FBS was added to each well and cells incubated for a total of 72 hours under standard conditions. Cells were then harvested by trypsinization and used as described for downstream immunoblotting or drug sensitivity assays.

### RNAseq analysis of isogenically manipulated EAC cells treated with LCI133

RNAseq analysis of isogenic PTEN shRNA LOF HEC1-B EAC cells treated with LCI133. Total RNA from DMSO or LCI133 treated control shRNA or PTEN shRNA cells were sequenced by using Illumina GAIIX sequencer. Next, raw paired end FASTQ files were assessed for sequencing quality using FastQC (v0.12.1) and subsequently trimmed to remove adapter sequences and low quality bases with Trim Galore (v0.6.10). The trimmed reads were aligned to the human reference genome (GRCh38) using STAR (v2.7.11b), followed by sorting, indexing, and deduplication. Gene level quantification was performed using Salmon (v1.10.3). The resulting count matrices were used for filtering low expression genes, performing sample to sample normalization, and identifying differentially expressed genes (adjusted P value < 0.0001 and fold change > 2). Differential expression analysis and volcano plot generation were conducted using the DESeq2 and EnhancedVolcano packages in R (v4.5.2). Biological pathway annotation was carried out using hallmark gene sets via the Gene Set Enrichment Analysis (GSEA) tool (v4.4.0).

### Animal studies and histopathology

All procedures involving animals were approved by the Atrium Health Wake Forest Baptist Institutional Animal Care and Use Committee (IACUC). 1 x 10^6^ HEC1-B PTEN shRNA KD vs control shRNA were injected subcutaneously into the right flank of NSG mice in 100-μL RPMI-1640 (ATCC) (NOD.Cg-Prkdcscid Il2rgtm1Wjl/SzJ (NSG, Strain #005557, The Jackson Laboratory), respectively, using a 25-G syringe needle. Tumor dimensions were recorded regularly using digital Vernier caliper. Tumor volume was measured using the following formula: volume = 0.5 × length × (width)^2^. Once tumors reached approximately 50-75 mm3, AN3 CA mice were randomized into two groups (n = 10 mice per group) and HEC-1-B PTEN shRNA KD tumor bearing mice were randomized into two groups and began QD treatment with vehicle (5% DMSO + 35% PEG-400 + 5% cremephor + 55% sterile molecular-grade H_2_O) or LCI133 (40 mg/kg) administered by IP injection. Mice in each group were treated daily for 3 weeks or until humane endpoint was reached (tumor >2000 mm3 or body condition score ≤ 2). Animals were sacrificed by CO2 asphyxiation followed by confirmatory cervical dislocation. Tumor, spleen, liver, and kidney were collected during necropsy and preserved appropriately for downstream protein, RNA, and histopathologic analysis. For histopathology and immunohistochemistry, tissue sections were fixed in 4% paraformaldehyde. After fixation for a minimum of 24 hours, tissues underwent routine processing, were embedded in paraffin, sectioned at 4 μm, stained with either hematoxylin and eosin (H&E) or primary antibodies for IHC as indicated and evaluated by light microscopy by Wake Forest University School of Medicine pathologist. Images were processed and edited using Aperio Digital Pathology Slide Scanner software (Leica Biosystems, Deer Park, IL).

### GEMM model for EAC PTEN^fl/fl^ x PR^cre+^

The GEMM model for spontaneous endometrial carcinoma was as described (14). It involves crossing the PTEN^fl/fl^ mice with the progesterone receptor promoter driving cre recombinase in the endometrium (14). Mice were genotypes for the PTEN floxed allele and cre-recombinase transgene to generate PTEN +/- and -/- mice. The PTEN^fl/fl^ PR^cre+^ mice develop EAC at 4 weeks of age. PTEN loss status was evaluated by WB analysis of PTEN, AKT and p-AKT (Figure 7). All homozygous mice develop EAC, detected at autopsy from 4-6 weeks and tumor burden was assessed by measure of the ratio of the mass of the entire excised female reproductive tract (FRT) divided by the animal weight (FRT/animal weight) (Fig. 7). PTEN^fl/fl^ x PR^cre+^ mice were treated with LCI133 starting at 4 weeks of age for 21 days at 40 mg/kg/dose/day via IP injection. FRTs from vehicle vs LCI133 treated mice were fixed in 4% paraformaldehyde for H&E, Ki67 staining or tissue for Western blot or RTPCR. During breeding and antitumor studies, all mice were housed under standard conditions with 12-hour light/dark cycle maintained at 65-75°F (∼18-23°C) with 40-60% humidity. To confirm pathology associated with female animals, individual Pten wildtype, haploinsufficient, and knockout mice were sacrificed at the indicated time point by CO2 asphyxiation followed by confirmatory cervical dislocation. Female reproductive tract (FRT) and spleen were excised and subjected to weighing and histopathologic assessment of tumor and splenocyte isolation and quantification, respectively.

### *Ex vivo* isolation, propagation, IC_50_ sensitivity to LCI133 and nRNA analysis of primary human EAC cells

Under IRB-00135161 informed consent, we obtained fresh EAC tumor tissue from EAC patients at the time of surgery. The freshly obtained primary EAC tumor tissue was dissociated into single cell suspension by using the BD Horizon Dri Tumor and Tissue Dissociation reagent (TTDR). In brief, the tissue was first finely minced using scalpels in a petri disc containing 5 mL of RPMI media. The minced tumor tissue was further mixed with 5-mL of warm TTDR reagent and incubated at 37° C for 30 minutes with mild but frequent agitation in a shaker incubator. After incubation 25 mL of 1% BSA/DPBS/2mM EDTA was added to the dissociated tissue followed by passing through a 70 µm cell strainer into a fresh tube. Subsequently, the tube was centrifuged at 250 x g for 8 minutes, and the pellet was resuspended in 2 mL of 1X RBC lysis buffer and incubated for 15 minutes at RT. Again, the pellet was washed with 40 mL of 1% BSA/DPBS/2mM EDTA solution and centrifuge at 250 x g for 8 mins. Further, pellet was resuspended in appropriate medium then cell count and cell seeding was performed for IC_50_ cell viability assays in the presence of absence of LCI133 or AZD4573 and Western blot for PTEN and p-AKT. Confocal IF microscopy was used to quantitate EU-labeled nRNA. Viability was assessed each day in culture to be greater than 95%. EAC cells were grown for 24-72 hours. Total nRNA was determined using EU labeling of cells for two hours at which time cells were treated for 20 min with DMSO or LCI133 at 1 uM concentration followed by quantitation of the EU positive cells, nRNA+ by fluorescence confocal microscopy in LCI133 treated vs DMSO control. The fluorescent signal was quantitated using the ImageJ software.

### Whole exome sequencing (WES) and bioinformatics analysis

Genomic DNA was isolated from frozen EAC tissues corresponding to all *ex vivo* evaluated patient EAC isolates using the AllPrep DNA/RNA kit (Qiagen, Germantown, MD, USA), as directed. DNA was quantified using the Qubit dsDNA BR Assay Kit (Thermo-Fisher, Waltham, MA, USA). RNA integrity was evaluated with an Agilent 2100 Bioanalyzer profile (Agilent, Santa Clara, CA, USA). Sequencing libraries were prepared using the Twist Human Core Exome Kit (Twist, South San Francisco, CA, USA). Prior to library preparation, DNA was fragmented before ligation to indexed adaptors. Amplified indexed libraries were hybridized to probes and captured using streptavidin magnetic beads. The resultant purified exome-enriched libraries were quantified using the Qubit dsDNA BR Assay Kit and qPCR to ensure optimum cluster densities on the sequencing flow cell. The quality of the libraries were assessed with an Agilent Technologies 2100 Bioanalyzer using a High Sensitivity DNA chip. The libraries were sequenced on an Illumina NextSeq 2000 for 200 cycles (Illumina, San Diego, CA, USA). The anticipated mean depth of coverage was greater than 200x with a uniformity of coverage greater than 99%. Short-read FASTQ files generated from WES were analyzed using nf-core/sarek v3.9.0^42^, Nextflow v26.04.6, and the GRCh38 reference genome. The workflow included read-quality control, preprocessing, alignment, coverage analysis, variant calling, annotation, and software-provenance tracking. Variants identified by Mutect2 in GATK v4.6.1.0 were annotated using VEP,SnpEff, and InterVar^43^ to evaluate their functional impacts and clinical pathogenicity Mutect2 calls were read into R with maftools^44^ and filtered to remove likely germline and artifactual variants: VEP MAX_AF >=0.1% (1% if COSMIC-flagged), variant allele fraction <5%, read depth <20x, FLAGS genes^44^, and sites called in >=4 of the 8 unrelated patients. This retained 1,490 of 3,033 calls.

## Statistical analysis

The mean was chosen as the representative center value for all graphs. The percentage of G2 arrest and the percentage of apoptosis were analyzed using ANOVA, where the percentage is the response variable, and treatment is the independent variable. The arcsine square root transformation was performed on the percentages for the variance homogeneity assumption of ANOVA. The qRT-PCR gene expression data were analyzed using linear mixed model with repeated measures, where gene expression level is the response variable; treatment, time and treatment × time interaction were the independent terms. Western blot data were analyzed by linear mixed model with repeated measures, with the protein level as the response variable and treatment as the independent variable. This model allows the evaluation of treatment effect within each experiment. For the qRT-PCR and Western blot data, each gene or protein was analyzed separately. QQ plot was used to assess the normality assumption of the model, and log transformation was applied when needed. For each model described above, the overall effect and the difference from pairwise comparisons were evaluated. Positive False Discovery Rate (pFDR) was used to correct P values for multiple testing. pFDR < 0.05 was considered statistically significant. All statistical analyses were performed using GraphPad Prism 9 (Boston, MA, USA).

