## Supplementary Data for "*In silico* engineered multitarget-directed ligands for the polypharmaceutical treatment of PTEN loss of function endometrial adenocarcinoma"

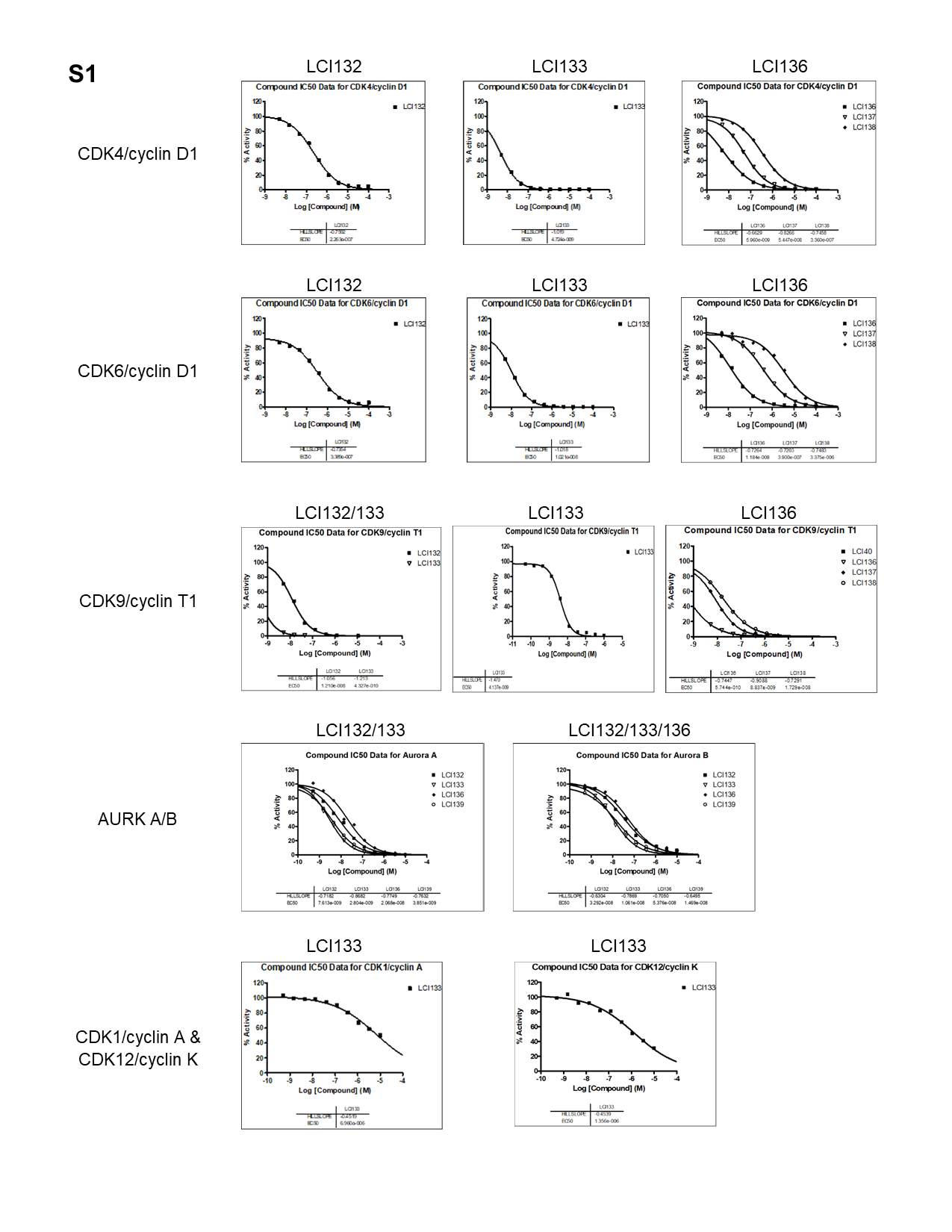

**Fig. S1 to support Fig. 1: LCI132, LCI133, and LCI132 are highly selective and potent multitarget small molecule inhibitors.** Cell free data, IC50 curves for LCI132, LCI133 and LCI136 against (*top to bottom*): CDK4/cyclin D1, CDK6/cyclinD1, CDK9/cyclinT1, AURKA and AURKB, and CDK1/cyclin A and CDK12/cyclin K using *in vitro* ATP-dependent kinase assays for each kinase target.

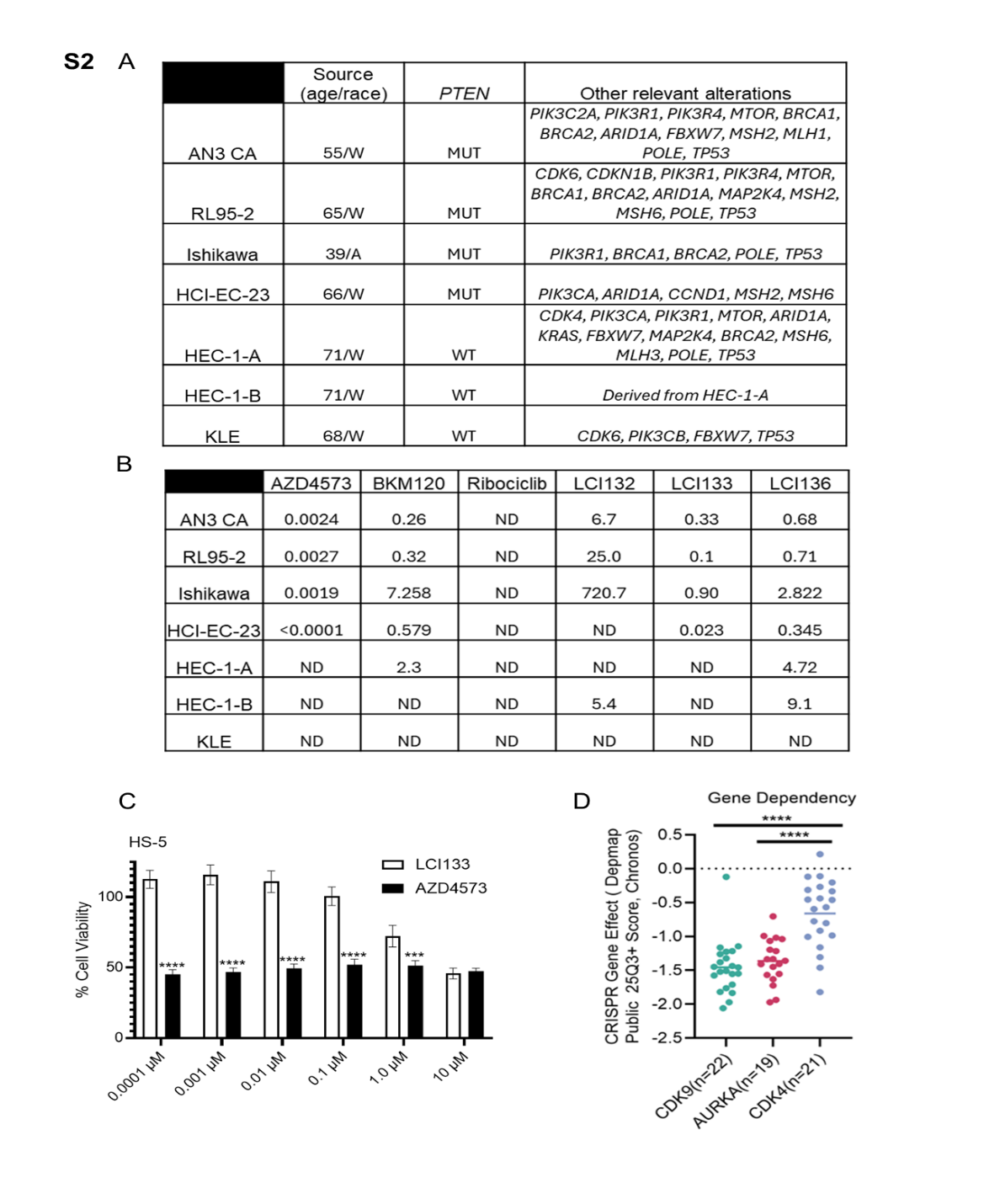

**Fig. S2 to support Fig. 2: PTEN mutational status of EAC cell lines, table of cell-based IC50 data for 48- hour treatment of EAC cell lines with AZD4573, BKM120, ribociclib, LCI132, LCI133 and LCI136, and relative stromal toxicity of LCI133 versus AZD4573. Fig. S2 to support Fig. 2: PTEN mutational status of EAC cell lines, table of cell-based IC_50_ data for 48-hour treatment of EAC cell lines with AZD4573, BKM120, ribociclib, LCI132, LCI133 and LCI136, and relative stromal toxicity of LCI133 versus AZD4573.** (**A**) Reported PTEN status (mutant, MUT; wildtype, WT) and other oncogenic alterations determined in each EAC cell line under study herein, sourced from the Dependency Map (DepMap) portal. **B**, EAC cell lines shown in Fig. 2 were treated with increasing logarithmic concentrations of the indicated inhibitors for 48-hours to obtain an IC_50_ curve for each compound for each respective cell line. Tabulation of IC_50_ values shown in Fig. 2B-H. **ND** indicates experiment performed but an acceptable IC_50_ curve was not determined as a non-linear curve to fit of the data. **C,** Comparison of AZD4375 vs LCI133 cytotoxicity to normal non-transformed HS5 bone marrow stromal cells following 48-hour treatment and viability assessment by CellTiter-Glo. Comparison AZD vs LCI133 where P value: **** p ≤ 0.0001, *** p ≤ 0.001. Data represent mean of triplicate values ± SEM. **D,** Means of dot plots depicting the CDK9, AURKA and CDK4 dependencies in EC cell lines by analyzing data from genome wide cancer genetic vulnerability CRISPR screen on DepMap. Number below shows the cell lines analyzed for each gene. A lower Chronos score indicates a higher likelihood that the gene of interest is essential in each cell line. A score of 0 indicates not essential. Data were analyzed by two-way ANOVA where P value**** p ≤ 0.0001.

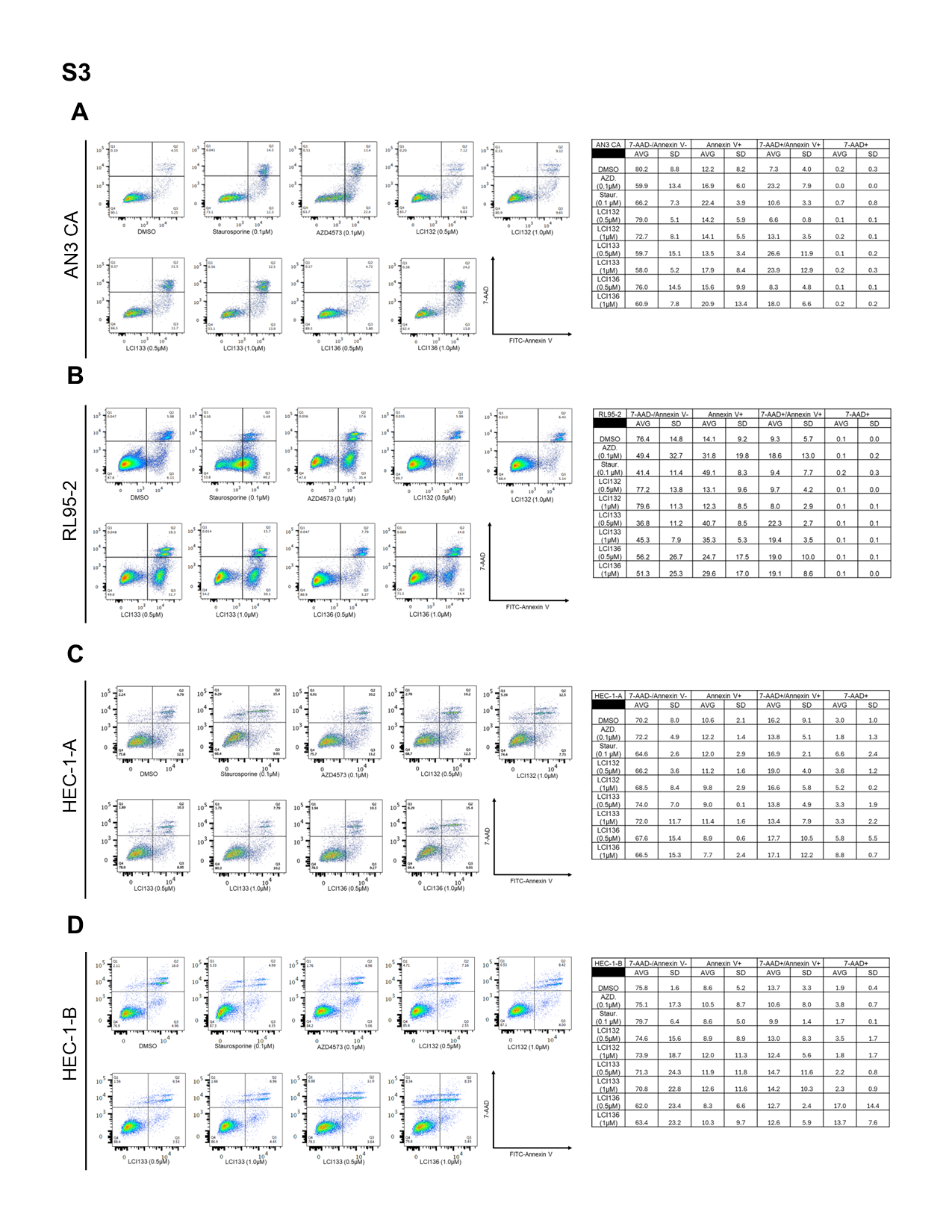

**Fig. S3 to support Fig. 3A-E: FACS plots and table of values for apoptosis. EAC cells are differentially sensitive to apoptosis induced by single agent and multitarget inhibitors. A-D,** Effect of single agent controls and LCI132, LCI133, and LCI136 on AN3-CA (**A**) RL95-2 (**B**) HEC1A (**C**) and HEC 1-B (**D**) EAC cell lines treated with the indicated concentration of inhibitor for 24-hours followed by flow cytometry assessment of FITC-Annexin V/7-AAD staining. Table shows the percentage of EAC cells in each phase of programmed cell death, representative of n = 3 replicates ± SD.

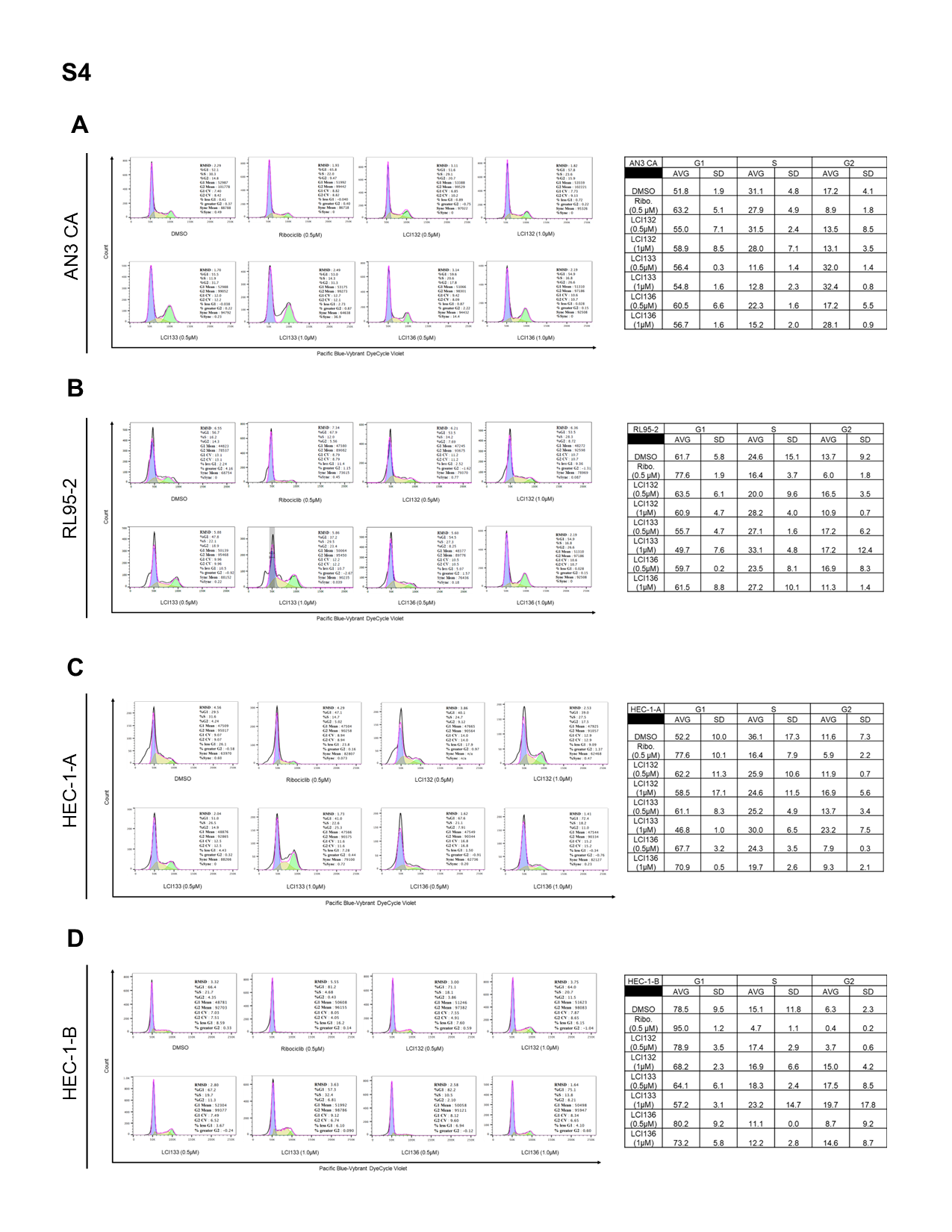

**Fig. S4 to support Fig. 3F-J: FACS plots and quantitative table of values (Fig.3F-J,), cell cycle. EAC cells are differentially sensitive to cell cycle arrest induced by single agent and multitarget inhibitors. A-D,** Effect of single agent controls and LCI132, LCI133, and LCI136 on AN3-CA (**A**) RL95-2 (**B**) HEC1A (**C**) and HEC 1-B (**D**) EAC cell lines treated with the indicated concentration of inhibitor for 24-hours followed by flow cytometry assessment of DNA binding dye staining. Table shows the percentage of EAC cells in each phase of the cell cycle as determined by dye MFI and curve fit model (FloJo), representative of n = 3 replicates ± SD.

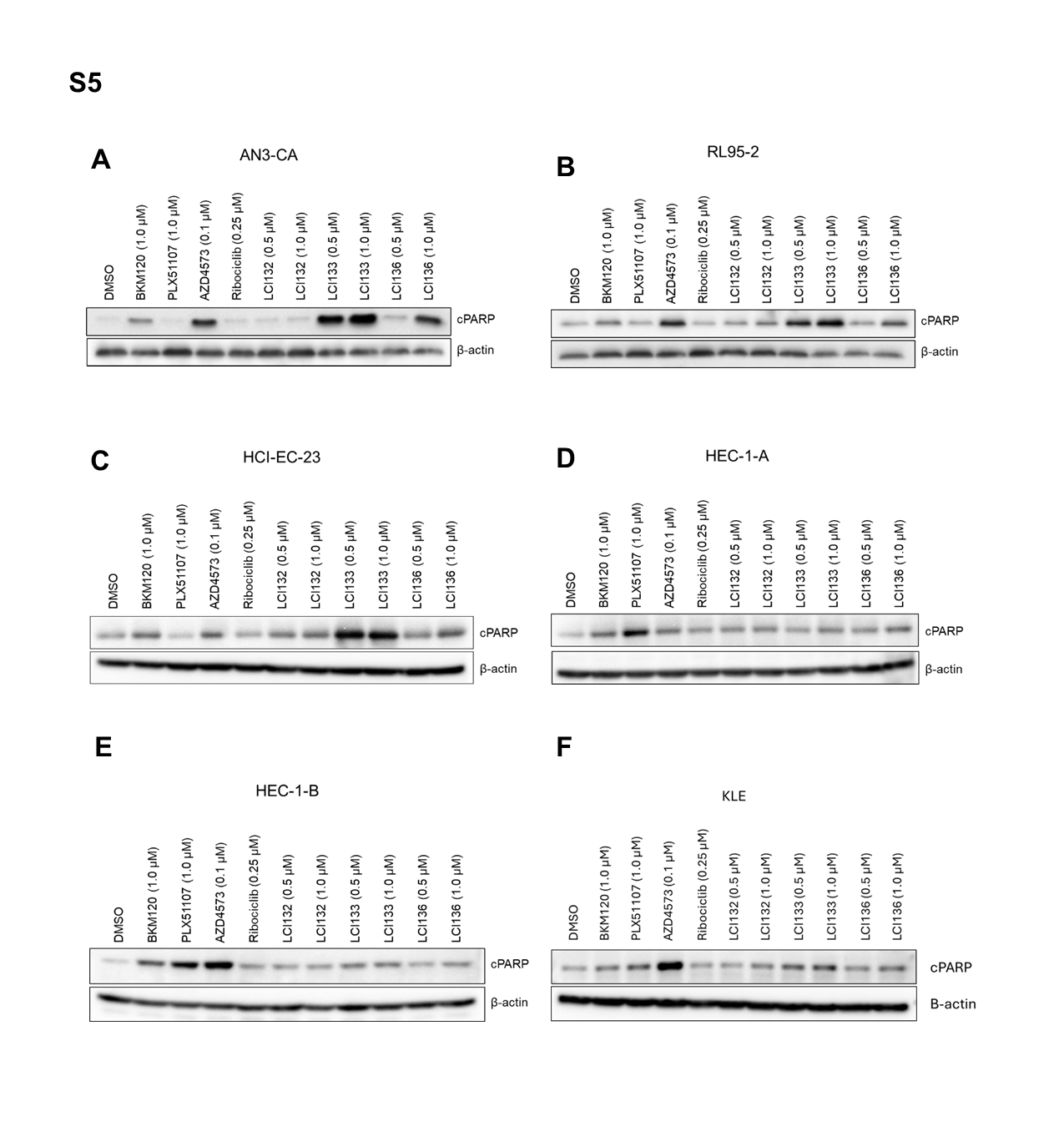

**Fig. S5 to support Fig. 3A-E: EAC cells are differentially sensitive to apoptosis (cleaved PARP, cPARP) induced by single agent and multitarget inhibitors. A-F,** Immunoblots showing effect of 24-hour drug treatment on apoptosis as determined by PARP cleavage in PTEN-mutant AN3 CA (**A**), RL95-2 (**B**), HCI -EC- 23 (**C**) vs PTEN wildtype HEC-1-A (**D**), HEC-1-B cells (**E**), KLE (**F**) EAC cell lines. AN3-CA, RL95-2, HCI-EC-23, HEC-1-A, HEC-1-B and KLE EAC cell lines were treated with the above labeled chemotype concentrations for 24 hours followed by detection of apoptosis by Western blot analysis for cPARP. β actin serves as loading control.

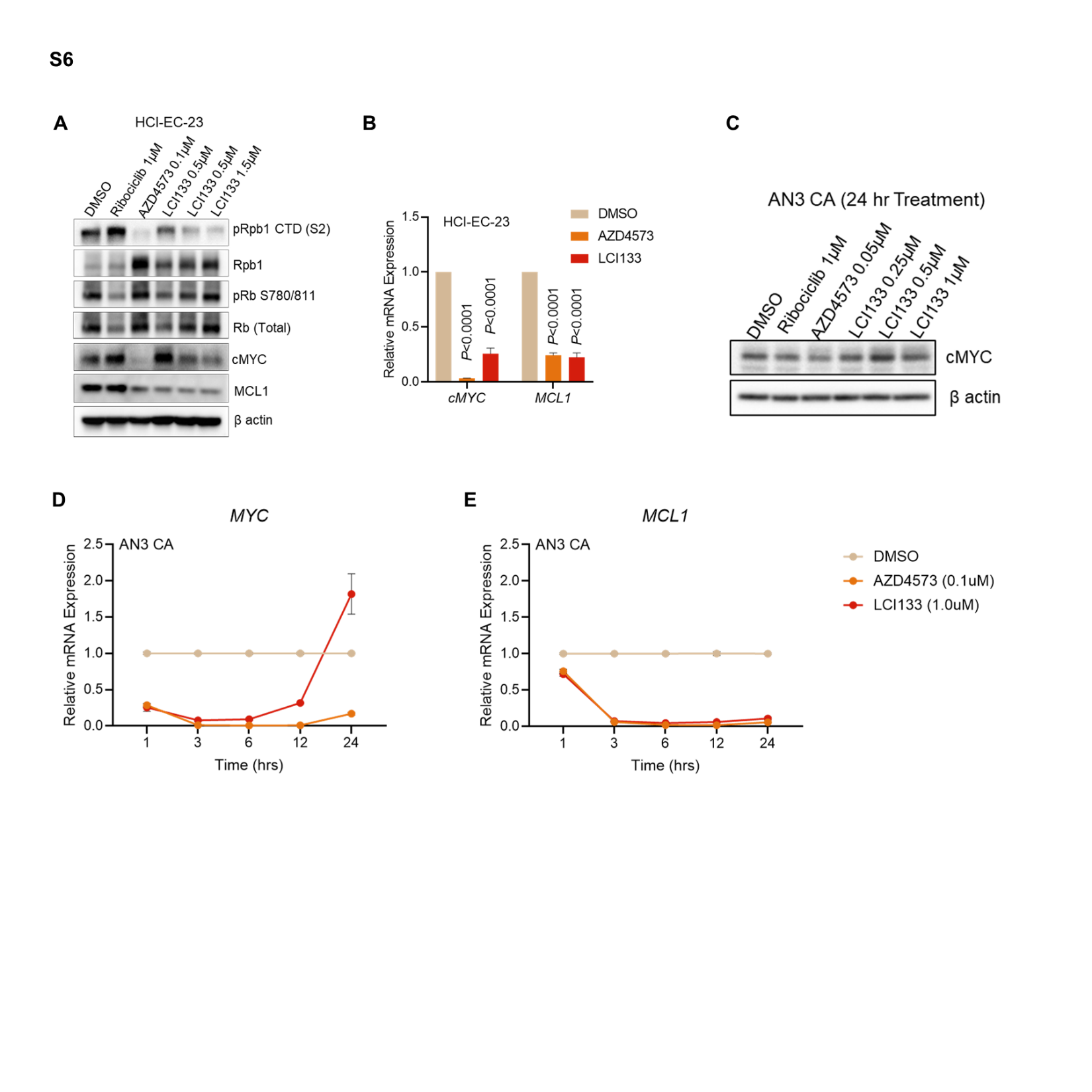
**Fig. S6 to support Fig. 4: LCI133 regulates *MYC* and *MCL1* transcription in PTEN-mutant EAC Cells via Pol II inhibition. A**, Western blot image showing the effect of LCI133 and the positive control single agent inhibitors AZD4573 (CDK9) and ribociclib (CDK4/6) on RNA polymerase II (Rpb1), phosphorylated RNAP II at serine 2 in the C-terminal domain (p-Pol II CTD Ser 2), pRb at S780/T 811, total Rb, MYC and *MCL1* in PTEN Mut EAC cell line HCI-EC-23 following 6-hour treatment. β actin served as loading control. **B**, qRT-PCR data showing the relative mRNA expression of *MYC* and *MCL*1 in HCI-EC-23 EAC cell line treated with DMSO control, AZD4573 (0.1µM), LCI133 (1 µM) for 2-hours followed. **C-E**, AN3 CA cells were treated at time zero with the indicated concentration of DMSO or experimental compounds for up to 24 hours to compare the kinetic effects of single agent inhibitors or LCI133 on MYC protein (**C**) and *MYC* (**D**) and *MCL1* (**E**) gene expression at different time points after exposure to each compound. β actin served as loading control. Bars represent the mean from n = 3 experiments ± SEM.

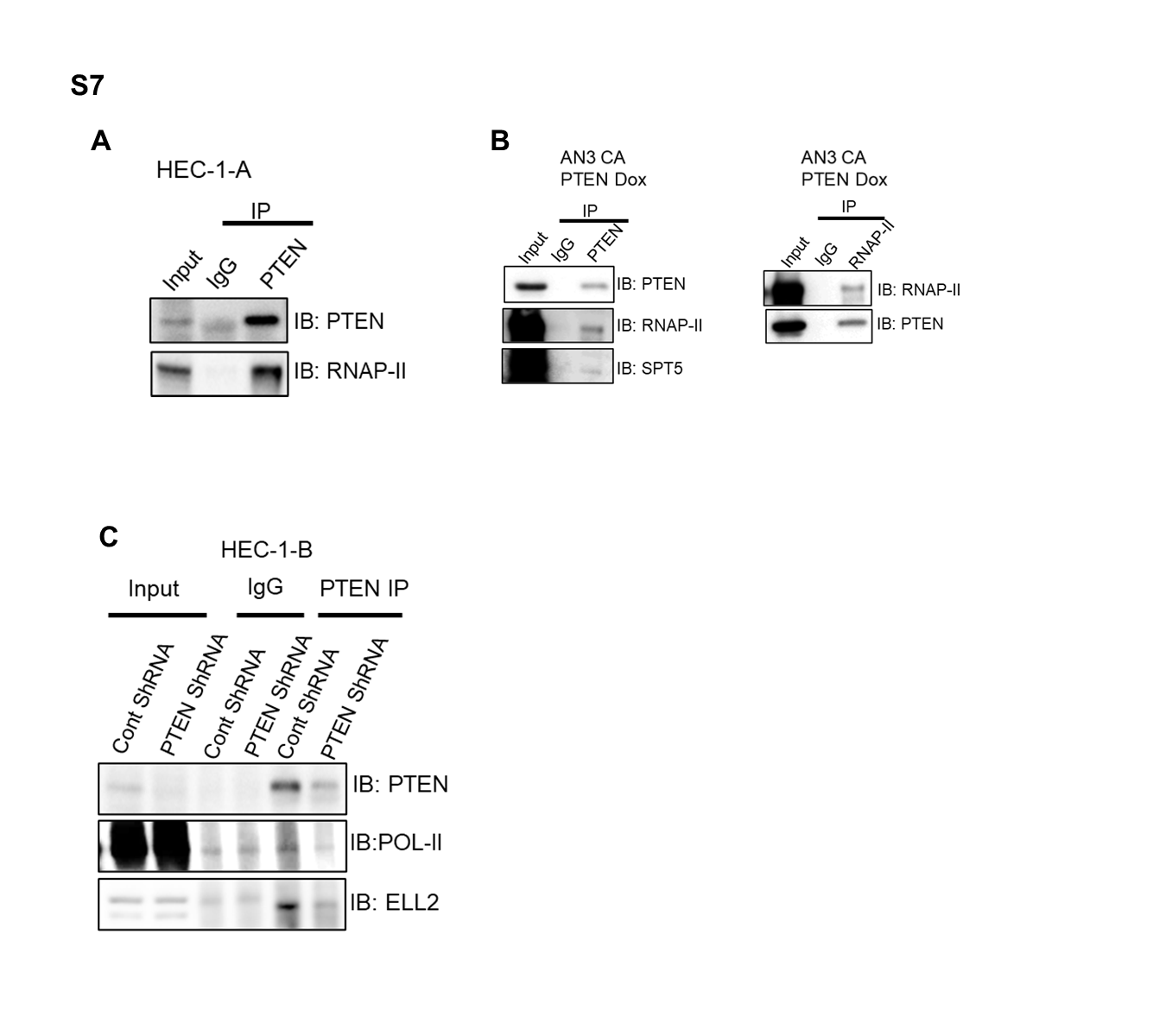

**Fig. S7 to support for Fig. 5: PTEN interacts with Pol II and super elongation complex (SEC) proteins. A,** Co-IP of PTEN and Pol II in PTEN wild type EAC cell line HEC-1-A. **B,** Co-IP and reciprocal Co-IP of PTEN or RNAP-II in Dox inducible PTEN in PTEN null AN3 CA (AN3 CA PTEN Dox) followed by Western blot analysis demonstrates a protein-protein interaction between Pol II and PTEN that is PTEN dependent. (**C**) PTEN wildtype HEC-1-B EAC cells were treated with PTEN shRNA or control shRNA to develop stable shRNA expressing cell lines. Immunoprecipitation of HEC-1-B-ShRNA cells with PTEN antibody followed by Western blot for Pol II and/or SEC protein ELL2. Representative Western blots where Input served as loading control.

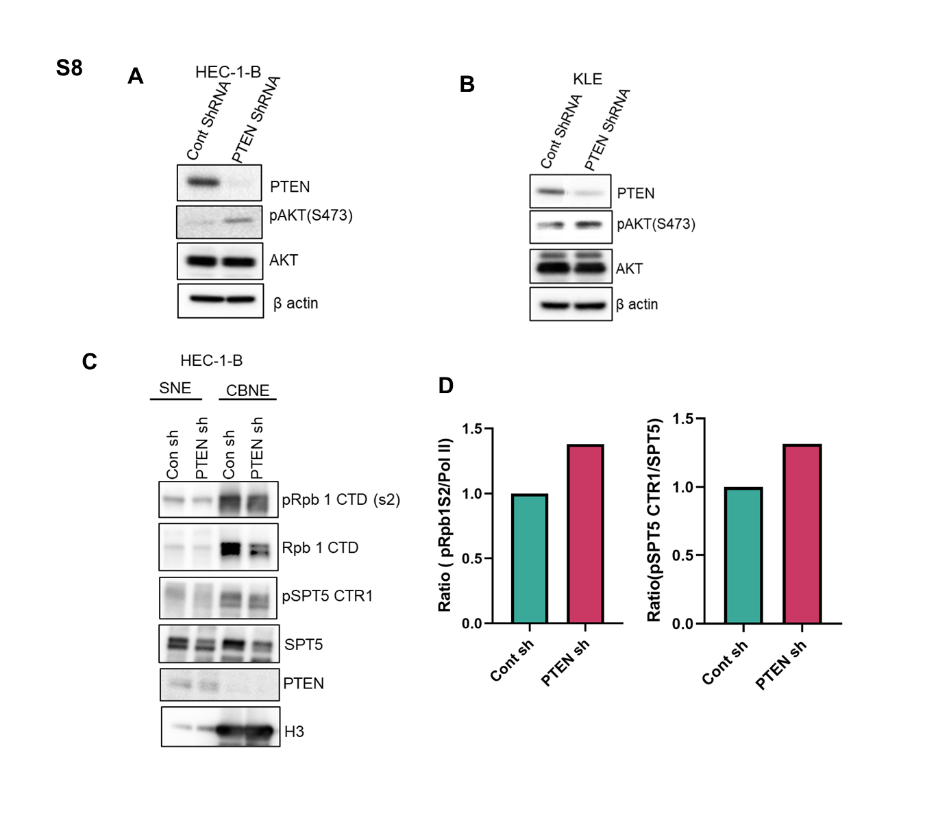

**Fig. S8 to support Fig. 5: shRNA mediated stable knockdown of PTEN in PTEN WT HEC-1-B and KLE EAC cells**. **A** and **B**, Western blot analysis of PTEN, p-AKT (S473), and total AKT in Control and PTEN shRNA transduced HEC-1-B (**A**) and KLE (**B**) EAC cells. β actin served as loading control. **C,** Westernblot analysis of pRpb1 S2, Rpb1, pSPT5 CTR1, SPT5, PTEN and H3 in soluble nuclear extract (SNE) and chromatin-bound nuclear extract (CBNE) from HEC-1-B cells stably expressing cont shRNA and PTEN shRNA. **D,** Bar graphs represent ratio calculated between phosphorylated Pol II s2/ total Pol II and pSPT5 CTR1/total SPT5 and quantified using ImageJ software (n=1 biological independent experiment)

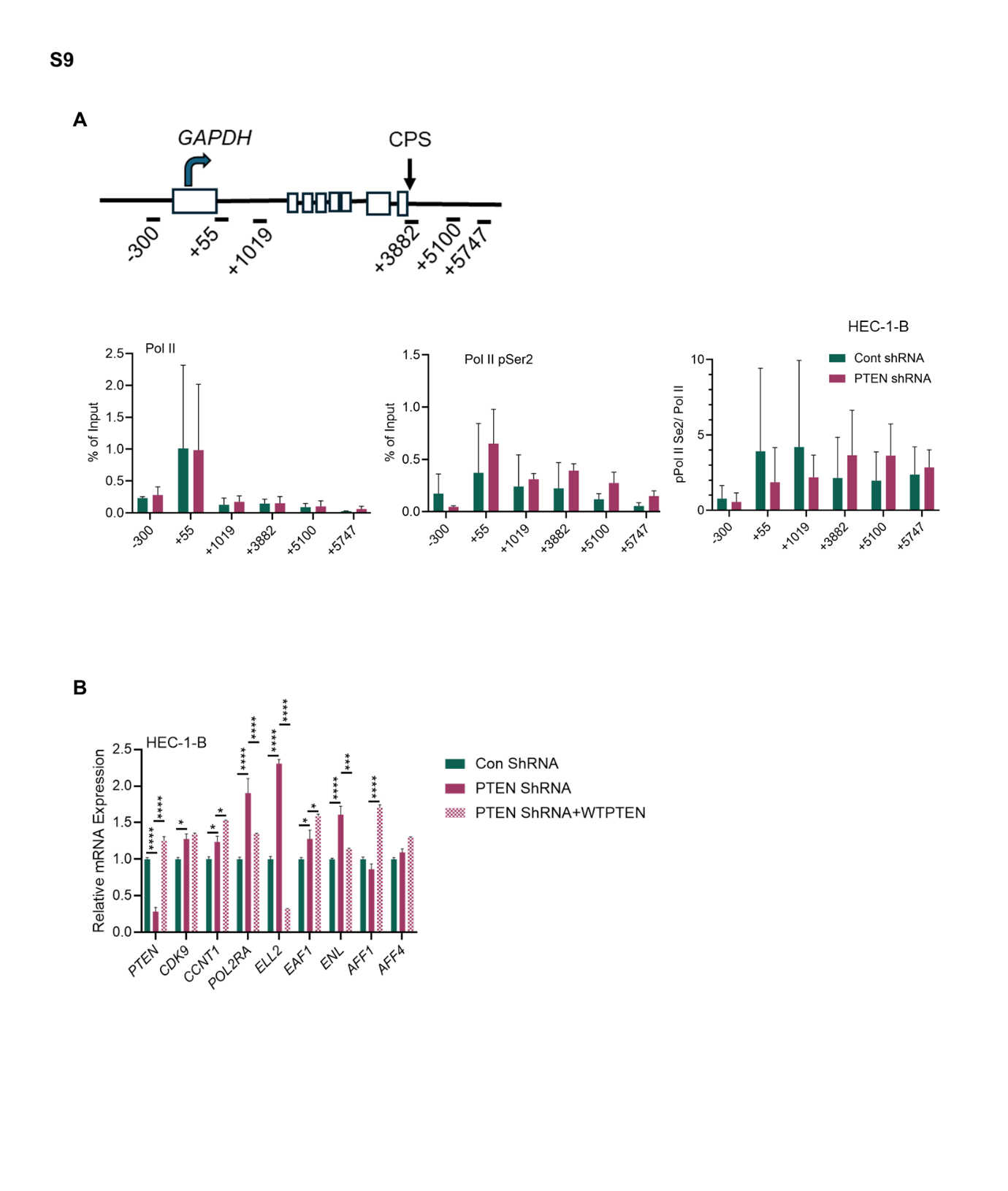

**Fig. S9 to support Fig. 5: PTEN knockdown increases Pol II, Pol II Ser2 occupancies on chromatin and alters key SEC gene expression in EAC cells. A**, Schematic diagram of the GAPDH promoter regions under study including TSS, gene body, and termination zone (*upper*). ChIP qPCR analysis of of Pol-II, Pol II Ser2 on GAPDH promoter and gene bodies in PTEN knockdown cells (*lower*). Error bars indicate ± SD of two biological replicates. **B**, qRT-PCR analysis of super elongation complex (SEC) factors in isogenic HEC-1-B control shRNA, PTEN shRNA and PTEN shRNA+wtPTEN cells under resting conditions. Error bars represent mean ± S.E.M from triplicate experiments. Data were analyzed using a two-way ANOVA where P value, **** p ≤ 0.0001, ***p ≤ 0.001, **p ≤ 0.01 or * p ≤ 0.05 denotes a significant difference in Con shRNA vs PTEN shRNA or PTEN shRNA vs PTEN ShRNA+wtPTEN.

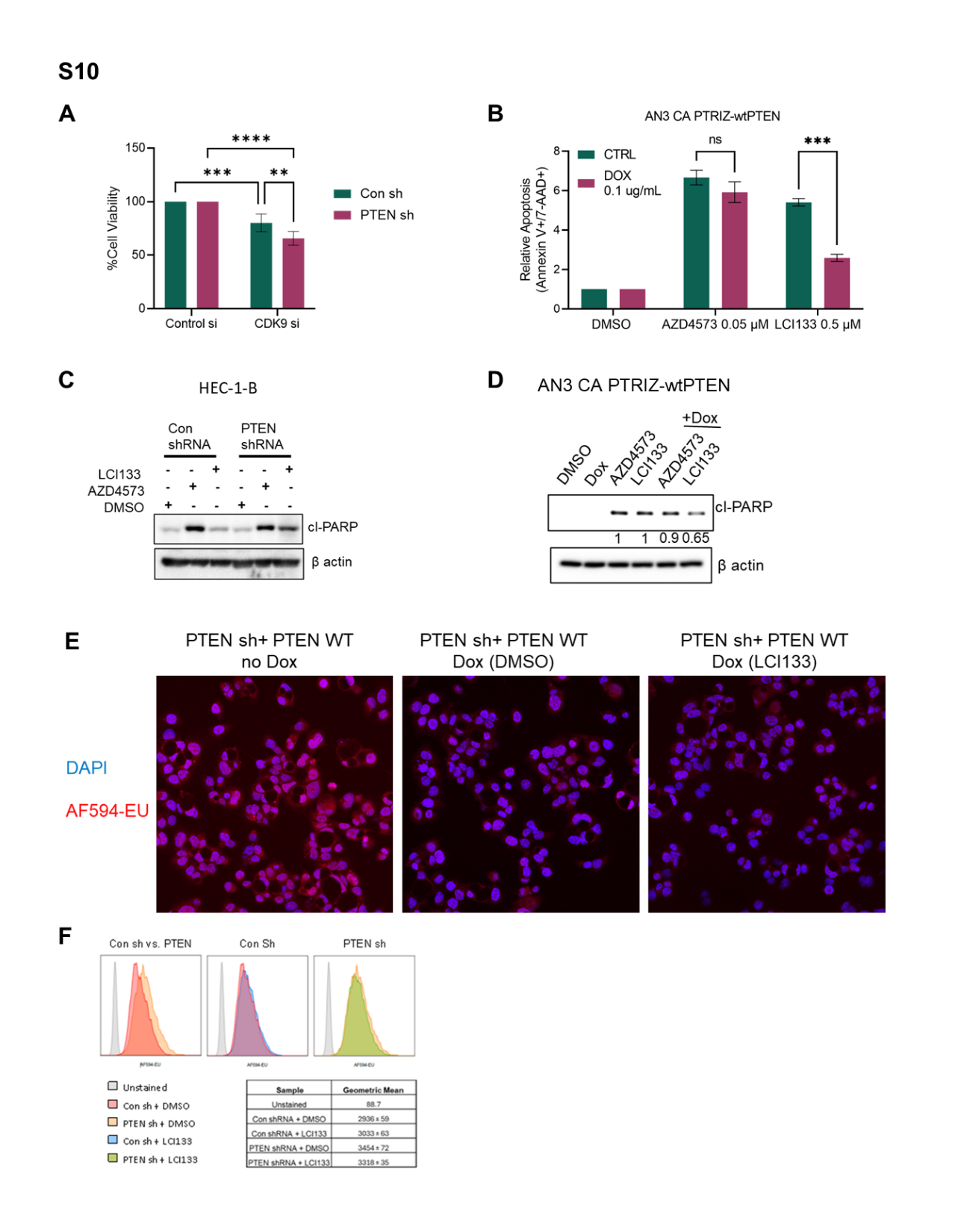

**Fig. S10 to support Fig. 6: PTEN knockdown or expression in EAC cells affects sensitivity to LCI133 and AZD4573 *in vitro* and levels of global nRNA. A**, AN3 CA cells were pretreated with DMSO or dox (1μg/ml) for 24-hours followed by treatment with DMSO control, AZD4573 (0.1 μM), or LCI133 (1.0 μM) and assessed for apoptosis by flow cytometry staining of FITC-Annexin V/7-AAD Data represent the mean ± SEM of duplicate experiments where P value, *** p ≤ 0.001. **B**, Western blot analysis of cl-PARP (cl-PARP) in control and PTEN shRNA HEC-1-B EAC cells treated with DMSO, AZD4573 (0.1 μM), or LCI133 (1.0 μM) for 48 hours. β- actin serves as loading control. **C**, Western blot analysis of cl-PARP in control and Dox treated AN3 CA PTEN Dox cells treated with DMSO, AZD4573 (0.1 μM), or LCI133 (1.0 μM) for 24 hours. Numbers below the blot show the quantification of band intensities after normalization to respective β- actin loading control in comparison with no DOX control samples using ImageJ software. **D**, Re-expression of PTEN WT in PTEN shRNA HEC-1-B cells treated with Dox

or no Dox (1µg/ml) for 24 hours, synchronized with 5,6-dichloro-1-β-d-ribofuranosylbenzimidazole (DRB; 100 µM) for 3 hours then washed, resuspension in fresh medium, treated with DMSO or LCI133 (1 µM), 20-minute EU incorporation, and click chemistry-mediated detection of EU-nRNA. Representative confocal microscopy images show nuclear DAPI (blue) and EU-nRNA (red). **E** and **F,** HEC-1-B-Con- and PTEN-ShRNA cells were synchronized with 5,6-dichloro-1-β-d-ribofuranosylbenzimidazole (DRB; 100 µM) for 3 hours followed by washout, resuspension in fresh medium, treatment with DMSO or LCI133 (1 µM), 20-minute EU incorporation, and click chemistry-mediated detection of EU-nRNA. Representative flow cytometry histograms (*upper*) show geometric mean distribution of AF-594-EU MFI in Con shRNA vs PTEN shRNA cells treated as above. Data are representative of technical duplicates per sample with average geometric mean ± SEM quantified in table (*lower*).

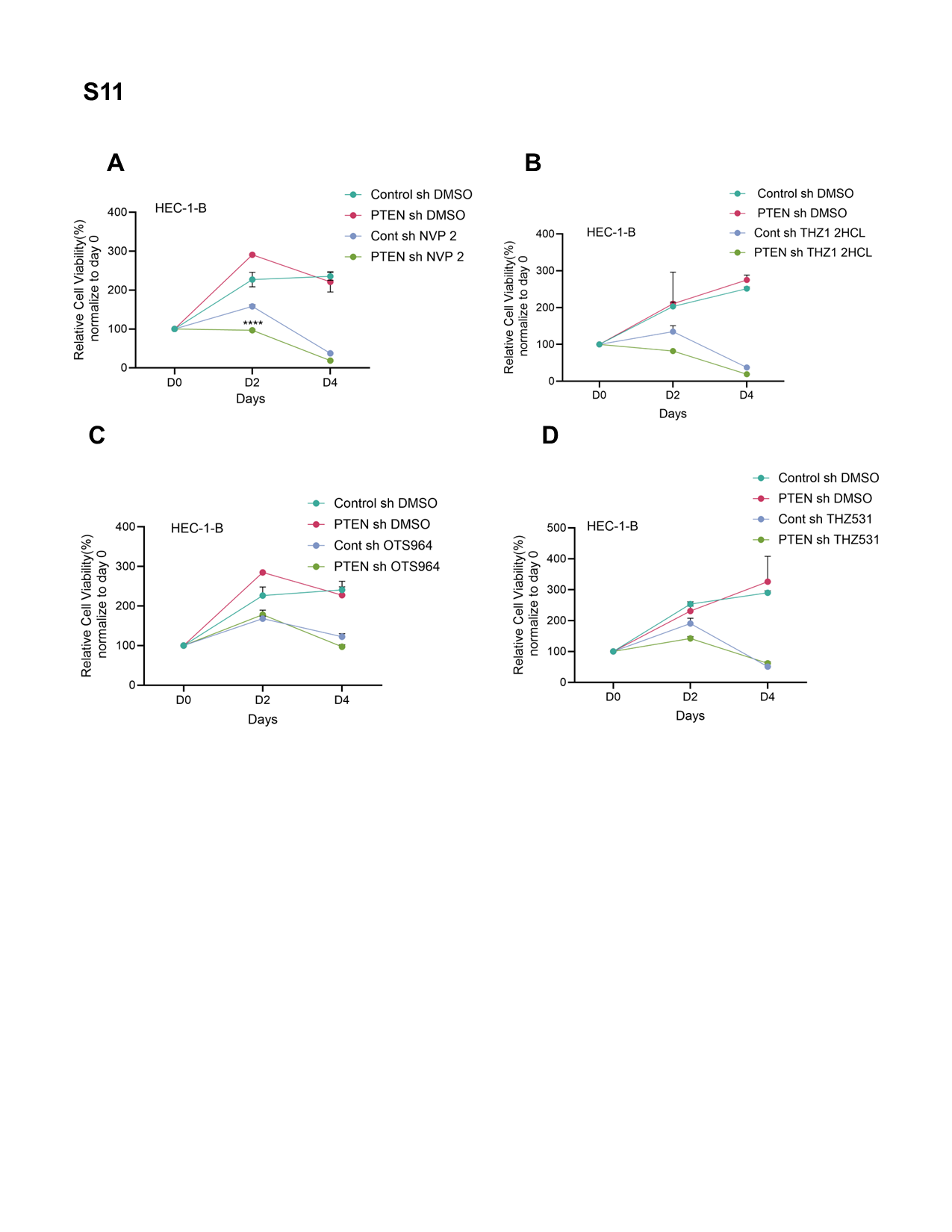

**Figure S11 to support Fig.6. Differential sensitivity of transcriptional inhibitors in PTEN knockdown EAC cells. A-D,** The HEC-1-B stably transduced control shRNA or PTEN shRNA cells were treated with specific CDK9i NVP2 (0.1 µM) (**A**), CDK7i (0.5 µM) (**B**), CDK11i OTS964 (0.25 µM) (**C** ), and CDK12i THZ531 (0.5 µM) (**D**) as indicated time. Cell viability was measured by using CellTiter-Glo assay reagents at each time point. Error bars represent ± S.E.M from triplicate experiments. Data were analyzed using a two-way ANOVA where P value, **** p ≤ 0.0001 denotes a significant difference in control sh vs PTEN sh.
